# Wildfire emitted particulate matter induces ovarian hyperandrogenism through aryl hydrocarbon receptor activation

**DOI:** 10.64898/2026.02.19.701615

**Authors:** Kuhelika Mali, Delong Zhang, Lila Bazina, Elena Abramova, Jiyang Zhang, Tingjie Zhan, Pawat Pattarawat, Kostas Moularas, Qiang Zhang, Audrey J. Gaskins, Andrew Gow, Philip Demokritou, Shuo Xiao

## Abstract

Wildfires have become more frequent and intense worldwide. Wildfire emitted particulate matter (WFPM) can be more toxic than urban background PM due to its greater content of nanoscale size (WFPM_0.1_) and presence of more polar organic compounds, including polycyclic aromatic hydrocarbons (PAHs). While exposure to WFPM has been linked to cardiovascular and respiratory diseases, its impact on female reproduction remains elusive. Here, we used an *in vivo* mouse intratracheal exposure model and a 3D ovarian follicle culture system, together with molecular, transcriptomic, and computational approaches, to examine the female reproductive effects of lab-synthesized (LS-WFPM_0.1_) and real-world Canadian WFPM_0.1_ (C-WFPM_0.1_), collected from the New York City and New Jersey metropolitan area during the June 2023 wildfire events. Intratracheal exposure to environmentally relevant dose of LS-WFPM_0.1_ disrupted mouse estrous cycles and elevated serum concentrations of estradiol and testosterone. RT-qPCR and single-follicle RNA-sequencing (RNA-seq) analysis revealed altered steroidogenic genes, transcriptomic changes, and activation of aryl hydrocarbon receptor (AhR) in antral follicles from mice treated with LS-WFPM_0.1_. LS-WFPM_0.1_ consistently increased testosterone secretion and stimulated genes related to androgen synthesis and AhR *in vitro*. Single-follicle and single-oocyte RNA-seq analysis identified differentially expressed genes related to inflammation in somatic cells and mitochondrial respiratory chain in oocytes. Both C-WFPM_0.1_ and benzo[a]pyrene, a high-molecular-weight PAH, reproduced these ovarian defects. Mechanistically, AhR inhibition reversed hyperandrogenism induced by WFPM_0.1_. Together, our findings suggest that WFPM_0.1_, an increasingly pervasive environmental exposure, adversely impacts female reproductive functions by disrupting ovarian steroidogenesis and inducing hyperandrogenism through AhR activation, highlighting an urgent unmet need for further mechanistic studies and epidemiological investigations to define the reproductive risks of wildfire smoke exposure in human populations.

## Introduction

Wildfires have been rapidly increasing in both frequency and intensity over the past decade, and this trend is only projected to continue in the coming years [1–6]. One major health concern of wildfires is the emitted particulate matter (PM). Ambient PM is classified into inhalable PM (< 10 μm or PM_10_), fine PM (< 2.5 μm or PM_2.5_), and ultrafine PM (< 100 nm or PM_0.1_). Wildfire emitted PM is primarily in the PM_0.1_ size range [7–9] and contains primarily organic compounds, including polycyclic aromatic hydrocarbons (PAHs). Approximately 70% of the California population in the United States experienced > 100 days of unhealthy air quality related to wildfires [10]. During the 2020 California wildfires, the daily mean PM_2.5_ reached 350-500 μg/m^3^ in the San Francisco Bay Area, far above the 24-hour limit of 35 μg/m^3^ [11]. The most recent wildfires in Los Angeles (LA) in January 2025 also generated average daily PM_2.5_ > 300 μg/m^3^ [12]. Wildfires are also spreading to historically less affected areas. In June 2023, New York City and New Jersey experienced their worst air pollution on record, with PM_2.5_ skyrocketing to > 400 μg/m^3^ due to the smoke emitted from the wildfires in Quebec, Canada, hundreds of miles away [7, 13, 14]. Another wildfire event in Alberta, Canada, in May 2024 recorded the ground PM_2.5_ concentrations as high as 949 μg/m^3^ [15]. Wildfires also impact other continents, including Australia, South America, Europe, Africa, and Asia [16–19]. Worldwide, > 2 billion population are exposed to ∼ 10 days of substantial wildfire pollution per year [20].

Epidemiological studies revealed that compared to urban background PM_2.5,_ which are mostly generated from the fossil fuel combustion, similar exposure levels of wildfire PM_2.5_ (WFPM_2.5_) are associated with higher risks of respiratory [21–24], cardiovascular [24–26], and mental health disorders [27], as well as a higher all-cause mortality [28, 29]. Toxicological studies using *in vitro* and *in vivo* models reported that WFPM_2.5_ exposure increases the expression of genes related to xenobiotic metabolism [30], cytotoxicity [31], and inflammation in bronchial epithelial cells [32], elevates reactive oxygen species (ROS) in bronchoalveolar lavage fluid (BALF) [14], and decreases the viability of alveolar macrophages [33].

The toxic mechanisms of WFPM_2.5_ might be due to their higher content of nano-sized or PM_0.1_ and more polar organic compounds with high oxidative potential, including PAHs [7–9, 34–39]. The chemical composition of WFPM is complex and highly variable, depending on fuel type, combustion conditions, and atmospheric transport, and may include heavy metals, plasticizers, flame retardants, industrial solvents, and PAHs generated from incomplete combustion [8, 9, 35–39]. Our recent publications revealed that the real-world WFPM collected from the New York City and New Jersey metropolitan area from the Canadian wildfires in June 2023 contains a significant amount of WFPM_0.1,_ which primarily consists of organic compounds (96%), and elemental analysis revealed potassium as the dominant element, along with other crustal metals and trace elements [7, 9, 14]. Organic speciation analysis identified high levels of PAHs in the Canadian WFPM_0.1_, including 8 out of 16 PAHs prioritized by the US Environmental Protection Agency (EPA) due to their persistence and toxicity [40, 41].

WFPM, particularly nanosized WFPM_0.1_, can distribute to the deepest part of the lungs, cross the alveolar epithelium, and enter blood circulation to accumulate in other organs, including those in the reproductive system [34, 42–45]. Existing studies have documented robust associations between higher exposure to all-source PM_2.5_ and a range of female reproductive dysfunctions [46–65], including diminished ovarian reserve [46–49], irregular menstrual cycles [50], longer time to pregnancy [51], higher risks of miscarriage [52–55], decreased live birth rates following infertility treatment [56–62], and lower county-level fertility rates [63–65]. However, there are currently limited mechanistic studies regarding the female reproductive impacts of WFPM. Given the unique and complex physicochemical properties of WFPM and its impact on air quality during wildfire accidents across the globe, there is an urgent unmet need to investigate the effects of WFPM on female reproductive health and elucidate mechanisms involved. In the current study, we used a human-relevant *in vivo* mouse model and a 3D *in vitro* ovarian follicle culture system, together with molecular biology, transcriptomic, and computational approaches, to investigate the ovarian impacts of WFPM_0.1_. We studied both lab-synthesized (LS)-WFPM_0.1_ and real-world WFPM_0.1_ collected from the central New York City and New Jersey area during the Canadian wildfire catastrophic events in June 2023.

## Materials and Methods

### Animals

CD-1 female mice used for *in vivo* WFPM_0.1_ exposure experiments were housed in polypropylene cages in the Animal Care Facility of Research Tower at Rutgers University’s Busch campus. Food and water were given *ad libitum* while housing them in a temperature and humidity-controlled room with 12/12-hour light/dark cycles. Animals were maintained by certified staff according to the NIH Guideline for the Care and Use of Laboratory Animals. All animal experiments were approved by the Institutional Animal Care and Use Committee (IACUC) at Rutgers University.

### Generation of LS-WFPM_0.1_

LS-WFPM_0.1_ were generated using a WildFire Simulator (WiFS) as described in our previous studies [9, 66–69] and in Figure S1. WiFS allows the incineration of wood samples under controlled combustion conditions and the production and collection of released PM for physicochemical characterization and exposure studies. The operational parameters of the combustion conditions were configured to simulate a flaming combustion scenario during wildfires with the final temperature at 600°C with a heating rate of 20°C/min and ambient O_2_ concentration (20.9 vol% in air). The starting weight of commercial pinewood pellets (Pine Mountain Starter Stikk Fatwood, Royal Oak Enterprises LLC, Roswell, GA), a representative wood type commonly involved in wildfires in Canada and California, was fixed at 100 mg to ensure the reproducibility of the combustion atmosphere between replicate experiments.

### Collection and extraction of LS-WFPM_0.1_

The PM_0.1_ fraction of LS-WFPM, termed LS-WFPM_0.1_, was collected using the Harvard Compact Cascade Impactor (HCCI) [69] on pre-cleaned 47-mm-diameter polytetrafluoroethylene (PTFE) filters with 2-μm pores (Pall Corporation, Port Washington, NY, USA). Collected LS-WFPM_0.1_ were then extracted from the PTFE filters as described by Pal et al [70]. In brief, PTFE filters containing WFPM_0.1_ size fraction were placed in 50 mL beakers and immersed in 15 mL of 75% (v/v) ultrapure ethanol and subjected to bath sonication for 30-60 seconds. Extracted LS-WFPM_0.1_ particle suspensions in 75% ethanol were washed three times with 75 mL of cell culture grade water (Cytiva, USA) by rotary evaporation to produce an ethanol-free aqueous suspension of LS-WFPM_0.1_ for *in vivo* and *in vitro* exposures. The efficiency of the extraction of LS-WFPM_0.1_, calculated via gravimetric analysis of filters and dried suspensions, was ∼ 99%. To create a background or vehicle control for *in vitro* exposure studies, a sterile PTFE filter without PM was subjected to the same extraction procedure.

### Collection and extraction of the real-world Canadian WFPM_0.1_ (C-WFPM_0.1_)

The PM_0.1_ size fraction of C-WFPM was collected at Rutgers University’s Busch campus during the Canadian wildfire event in June 2023. Detailed WFPM_0.1_ collection and extraction methods are similar to those of LS-WFPM_0.1_ described above and have been reported in our previous publication [7].

### Real-time monitoring and physicochemical analysis of emitted WFPM_0.1_

During the combustion of pinewood pellets, emissions of LS-WFPM_0.1_ were monitored in real time using a Scanning Mobility Particle Sizer (SMPS, TSI Inc.) and an Aerodynamic Particle Sizer (APS, TSI Inc.) [3]. The physicochemical characterization and monitoring of LS-WFPM_0.1_ and C-WFPM_0.1_ were described in our recent publication [7, 8]. Both LS- and C-WFPM_0.1_ collected with HCCIs were characterized for their physicochemical property, including mass particle size distributions, organic and elemental carbon content, heavy metals, and PAHs, following previously established protocols [3, 6, 8–10]. Additional details are provided in the Supplementary Information.

### Endotoxin and microbiological sterility testing of LS-WFPM_0.1_ suspensions

To ensure the absence of biological contamination, endotoxin levels in the final LS- and C-WFPM_0.1_ aqueous suspensions were assessed as we previously reported [7]. In brief, the HEK-Blue™ LPS Detection Kit was used following both the manufacturer’s protocol and the approach described by Bazina et al. [14]. To further confirm microbiological sterility, all particle suspensions and corresponding vehicle controls were incubated in thioglycolate broth and observed for signs of microbial contamination. In parallel, bacterial and fungal growth were monitored over 14 days using conventional microbiological assays, consistent with methods applied in our previous studies [71, 72]. Following a 14-day incubation, agar plates exposed to LS- and C-WFPM_0.1_ and control vehicle aqueous suspensions showed no microbial growth. Additionally, endotoxin assessment of all samples revealed levels below the assay’s detection threshold of 0.0079 EU/mL, indicating the absence of detectable endotoxin. Further methodological details are available in the supplementary information.

### Mouse Intratracheal Instillation of LS-WFPM_0.1_

The collected LS-WFPM_0.1_ were suspended in HyPure endotoxin-free water for intratracheal administration. A tracheal catheter was assembled by inserting a 23-G needle into a 21-G transparent polyethylene tubing. Anesthesia was induced via intraperitoneal injection (i.p.) of short-acting barbiturate at 25 mg/kg using a 26G syringe needle. Anesthesia was confirmed by the absence of a foot withdrawal reflex. Mice were then placed in a supine position on a surgical platform, and the trachea was exposed by sequentially dissecting the skin, salivary glands, and sternohyoid muscle. An instillation 26G needle was used to carefully puncture the trachea between cartilage rings, after which the catheter was inserted approximately 0.5 cm into the tracheal lumen to avoid damaging the lungs and secured with a ligature to prevent backflow. Each mouse received an one-time intratracheal instillation of 443 µg LS-WFPM_0.1_ (8.86 µg/µL in 50 µL vehicle), equivalent to 75 days of WFPM exposure in humans as described below [73, 74] (Figure 1A). A sample size of 3-5 mice was included per dosing group.

**Figure 1.**
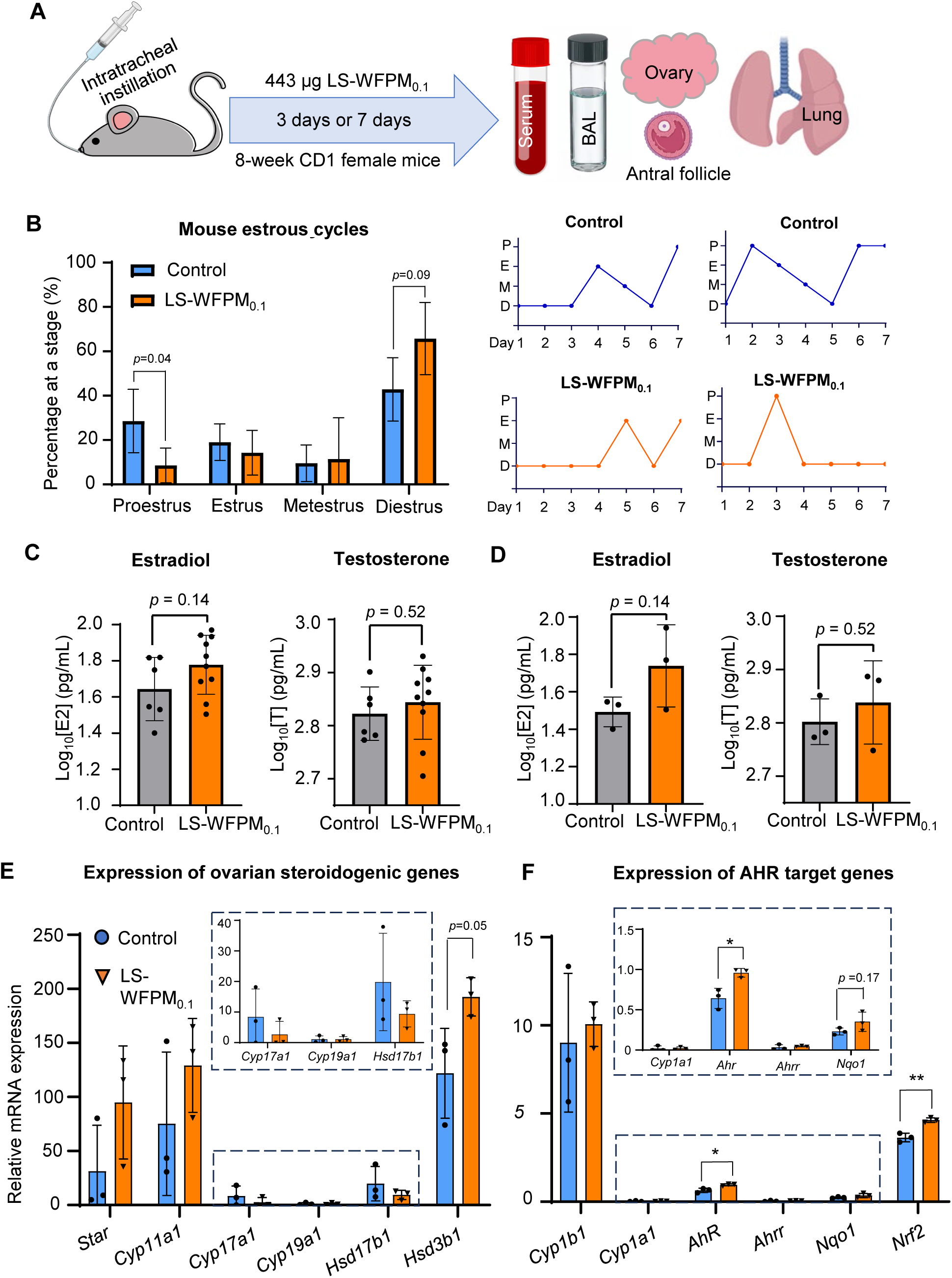
Effect of intratracheal instillation of 443 ug LS-WFPM_0.1_ on estrous cycle, serum hormone concentration and ovarian gene expression. (A) Schematic of intratracheal instillation in 8-week female CD-1 mice and sample collection after 3 days or 7 days. (B) Percent of days at each stage of mouse estrus cycle through a span of 7 days after exposure in control (N=3) and LS-WFPM_0.1_ treated (N=5) mice, with representative cycles from control (N=2) and LS-WFPM_0.1_ (N=2) mice. (C) Average Log_10_ serum concentration (pg/mL) of estradiol and testosterone from mice, including all stages of the estrous cycle, collected on day 3 and day 7 post-exposure in control (N=6) and (LS-WFPM_0.1_, N=10). (D) Average Log_10_ serum concentration (pg/mL) of estradiol and testosterone from mice only at proestrus, collected on day 3 and day 7 post-exposure in control (N=3) and (LS-WFPM_0.1_, N=3). (E-F) Relative mRNA expression of steroidogenic and AhR target genes in 3 pooled antral follicles derived from control (N=3) and LS-WFPM_0.1_ mice (N=3) examined by RT-qPCR. The mRNA expression levels were normalized by the expression of glyceraldehyde-3-phosphate dehydrogenase (*Gapdh*). Data was analyzed with Student’s t-test. Shown are mean ± standard deviation; \**p* 0.05.

Estimation of the WFPM instillation dose: The mass of WFPM_0.1_ deposited in the tracheobronchial and pulmonary regions of the human respiratory system was estimated to be 4.99 x10^-5^ µg/min/cm^2^ using the Multiple Path Particle Dosimetry (MPPD) model (V3.04) [75–77] based on aerosol characterizations during the June 2023 Canadian wildfire event and methods following previous publications [7, 78]. 165 µg/m^3^ of ambient mass concentration was used per day for 75 days of exposure, approximating long-term inhalation exposure during peak wildfire season. This analysis used the Yeh/Schum symmetric model [79] with a functional residual capacity of 3300 mL, a head volume of 50 mL, a nasal respiratory rate of 12 breaths/min, a tidal volume of 625 mL, and an inspiratory fraction of 0.5. Aerosol input parameters are provided in Table S11, and the effective density of WFPM_0.1_ was estimated based on measurements reported by Leskinen et al [78, 80]. These human deposition data were used to determine the one-time *in vivo* intratracheal instillation bolus dose of 443 µg (Figure 1A), while considering the mouse alveolar surface area of 82.2 cm^2^ [81].

### Blood and tissue collection

All blood and tissue collections were conducted under anesthesia following 3 days and 7 days of post-exposure. Anesthesia was performed via i.p. injection of a short-acting barbiturate at a dose of 25 mg/kg using a 26G needle. Successful anesthesia and euthanasia were confirmed by the absence of a foot withdrawal reflex. For blood collection, a small transverse skin incision was made just above the pelvis, followed by a midline incision extending cranially. The abdominal muscles were carefully opened along this line, and the ribcage was cut transversely along its natural curve, taking care to avoid damaging the liver. The intestines were gently moved to the left to expose the posterior vena cava. A sterile gauze pad was placed over the liver to reduce bleeding, and the needle was inserted into the posterior vena cava between the renal vessels and the iliac bifurcation. Blood was withdrawn slowly and gently while applying steady pressure to the liver to prevent hemorrhage and collected into a 2-mL Eppendorf tube. The blood was allowed to clot at room temperature for 30-60 minutes, then centrifuged at 1500 × g for 10 minutes. Serum was carefully collected from the upper layer and stored at −80°C for future use.

Following blood collection, lungs and ovaries were harvested. One ovary was transferred into dissection medium (L15 medium, Invitrogen, Carlsbad, CA) to isolate large antral follicles, which were then placed in lysis buffer for total RNA extraction. The second ovary from each mouse was fixed in 4% paraformaldehyde (PFA). One lung was fixed in 4% PFA, while the other was immediately snap-frozen.

### Bronchoalveolar lavage (BAL) collection

BAL samples were collected on day 7 post intratracheal instillation. To prepare the balanced salt solution for BAL collection, 100 μM EDTA was added to chelate divalent cations, along with protease inhibitors to prevent protein degradation. One mL of sterile balanced salt solution containing 100 μM EDTA was instilled through the catheter, followed by a gentle aspiration while massaging the thorax. 700-900 μL of fluid was recovered per lavage, and the procedure was repeated twice to obtain a sufficient sample volume. The cellular and acellular fractions were separated through centrifugation at 400 × g for 7 minutes at 4°C. The resulting cell pellet was resuspended in 200 μL of ACK lysing buffer for selective red blood cell lysis, incubated for 2 minutes at room temperature, diluted with cold PBS, and centrifuged again at 400 × g for 7 minutes at 4°C. The final cell pellet was resuspended in PBS for subsequent cytological or molecular studies. Cellular membrane integrity was evaluated by measuring lactate dehydrogenase (LDH, as the positive control indicating successful instillation) release in BAL fluid using the CyQUANT LDH Cytotoxicity Assay, following the manufacturer’s instructions. Further details are given in the Supplementary Information.

### 3D encapsulated *in vitro* follicle growth (eIVFG), exposure to LS-WFPM_0.1_ or C-WFPM_0.1_, follicle growth, and *in vitro* ovulation and luteinization

We have previously demonstrated that eIVFG faithfully recapitulates hallmarks of follicle development, hormone secretion, ovulation, and oocyte maturation, enabling a powerful *in vitro* tool to study the toxic effects and mechanisms of ovarian endocrine disrupting chemicals [82–90]. Herein, 16-day-old female CD-1 mice were euthanized in a CO_2_ chamber followed by cervical dislocation. Euthanized animals were dissected to collect ovaries on both sides, which were incubated in an enzymatic digestion solution containing L15 media (Invitrogen, Carlsbad, CA) with 30.8 µg/mL Liberase (Sigma-Aldrich) and 200 µg/mL DNase. The incubation was carried out for 25 minutes on a plate shaker in a 37°C, 5% CO_2_ incubator (Thermo Scientific). Secondary follicles with diameter range of 150-180 µM were isolated and encapsulated in 0.5% alginate hydrogel for encapsulation and *in vitro* culture as we previously described [82, 85, 90–92]. Briefly, follicles were encapsulated in 0.5% (w/v) alginate hydrogel (Sigma-Aldrich, St. Louis, MO) and were then individually cultured in 96-well plates. Each well contained a single follicle and 100 μL growth media consisting of 50% αMEM Glutamax (Thermo Fisher Scientific) and 50% F-12 Glutamax (Thermo Fisher Scientific) supplemented with 3 mg/mL bovine serum albumin (BSA; Sigma-Aldrich), 5 mIU/mL human recombinant follicle stimulating hormone (rFSH; gifted from Dr. Mary Zelinski from the Oregon Non-human Primate Research Center at the Oregon Health and Science University, Beaverton, OR, USA), 1 mg/mL bovine fetuin (Sigma-Aldrich), and 5 μg/mL insulin-transferrin-selenium (ITS, Sigma-Aldrich). Follicles were cultured for 4 days at 37 °C in a humidified environment of 5% CO_2_ in air, which allowed immature follicles to grow from the multi-layered secondary stage to the antral stage to reach maturation.

On day 4 of eIVFG, the grown early antral follicles were released from alginate by incubation in L15 media containing 1% FBS and 10 IU/mL alginate lyase (Sigma-Aldrich) at 37 °C for 10 minutes. Freed follicles were transferred to a 96-well round-bottom ultra-low-attachment plate (Corning®, Catalogue. No. 7007, Corning Life Sciences, Kennebunk, ME, USA) containing 100 μ L growth media with 10 mIU/ml recombinant FSH (rFSH, gifted by Organon, Jersey City, NJ, USA). Follicles were cultured from day 4 to day 6 in a 37°C, 5% CO2 incubator on a plate shaker (VWR standard orbital shaker, model 1000, Troemner, LLC, USA) to ensure uniform suspension of WFPM_0.1_ particles in the culture medium and prevent precipitation, thereby allowing consistent exposure throughout the *in vitro* follicle culture period. Follicles were randomly assigned to various experimental groups, with each group containing 10-15 follicles. The exposure groups included 0 (control, same amount of water), 1, 10, and 100 μg/mL concentrations of either LS-WFPM_0.1_ or C-WFPM_0.1_.

The *in vitro* doses used here are based on estimates from previous studies that equated 10 µg/mL of PM_2.5_ generated from wood combustion with an air pollution level of ∼50 µg/m³ [93–95]. Using this scaling, the treatments of 1, 10, and 100 µg/mL would correspond approximately to PM_2.5_ of 5, 50, and 500 µg/m³, respectively. Given the recorded June 2023 Canadian WFPM_2.5_ levels exceeding 400 µg/m³ in several regions, including New York City and parts of New Jersey [7, 13, 14], our highest *in vitro* dose (100 µg/mL) represents an environmentally relevant dose exposure to WFPM_0.1_, simulating severe but realistic wildfire conditions, while the lower doses correspond to lower exposure levels. It is also important to note that our study focuses on PM_0.1_, excluding the slightly larger PM_2.5-0.1_ fraction. Ultrafine particles are known to have higher surface area-to-mass ratios, enhanced cellular uptake, and typically elicit stronger oxidative and inflammatory responses than larger particles [96]. Therefore, our exposure concentration range of 1–100 µg/mL might capture the increased toxicological potency associated with ultrafine wildfire particles. Moreover, to translate environmental WFPM_0.1_ exposure into our eIVFG system, we used field measurements from the June 2023 Canadian wildfire event, which reported 13.8 ng/m^3^ total PAHs associated with 145.8 μg/m^3^ WFPM_0.1_ (≈ 0.0095% PAH by mass in WFPM_0.1_) [7]. Applying this measured PAH: WFPM_0.1_ mass ratio to our C- and LS-WFPM_0.1_ doses yields approximate total-PAH concentrations of 0.095, 0.95, and 9.5 ng/mL at 1, 10, and 100 µg/mL WFPM_0.1_, respectively. These values, and even higher concentrations, have been reported to fall within both experimental and modeled human biomonitoring ranges of PAH levels in blood [97–99]. Thus, the *in vitro* exposure range of 1-100 µg/mL WFPM_0.1_ brackets a realistic exposure window relevant to PAH levels in humans.

On day 6 of eIVFG, grown antral follicles were incubated in the ovulation induction media containing 50% αMEM Glutamax and 50% F-12 Glutamax supplemented with 10% FBS, and 1.5 IU/mL hCG (Sigma-Aldrich), and 10 mIU/mL rFSH. Ovulation was assessed at 14 hours post-hCG treatment using an Olympus inverted microscope with a 10x objective (Olympus Optical Co Ltd, Tokyo, Japan) with Tcapture imaging software (Tucsen, V5.1.1). A follicle was defined as “ruptured” when one side of the follicular wall was breached and defined as “unruptured” when the follicular wall was intact. Oocytes released from ruptured follicles were examined for the extrusion of the first polar body. Oocytes exhibiting a visible first polar body were classified as metaphase II (MII) oocytes, whereas oocytes that were either in the germinal vesicle stage (GV) or in the GV breakdown (GVBD) stage were classified as non-MII oocytes. The conditioned media from day 6 of culture were collected for subsequent hormone analysis using ELISA. Follicles after ovulation induction were continuously cultured in the ovulation induction media without rFSH for 48 hours to induce luteinization and progesterone secretion. The conditioned media were collected at 48 hours and stored at −20 °C for subsequent measurement of progesterone using ELISA.

During eIVFG, half of the follicle culture media was replaced every other day, and the conditioned media were used to measure steroid hormone concentrations by ELISA. Follicles were imaged at each media change every other day using the 10x objective of the Olympus inverted microscope for evaluating follicle survival and size. Follicle death was defined by the morphological changes of oocytes with shrinkage, irregular shape, and/or fragmentation, and/or morphological changes of follicles with darker and/or disintegrated somatic cell layers. The follicle size was determined by averaging 2 perpendicular measurements from one side to another side of the theca externa per follicle using the ImageJ software (v1.53; National Institutes of Health, NIH, Bethesda, MD).

### Hormone Assays

The concentrations of 17β-estradiol (E2), testosterone (T), and progesterone (P4) in the conditioned media were measured using ELISA kits (Cayman Chemical, Ann Arbor, MI, Research Resource Identifier (RRID): AB_2832924, catalogue: 501890 for E2 ELISA kit; RRID: AB_2811273, catalogue: 582601 for P4 ELISA kit; RRID: AB_2895148, catalogue: 582701 for T ELISA kit) according to the manufacturer’s instructions. The ELISA kits we used have also been widely used in hormone detection in many mouse-related studies [100, 101]. The mouse anti-rabbit IgG-precoated wells were incubated with standards, conditioned culture media, rabbit antiserum, and hormone-acetylcholinesterase (AChE) conjugate for 60-120 min. After the wells were washed with washing buffer three times, Ellman’s reagent was added and incubated at room temperature for 60-90 mins. The absorbance was measured using a BioTek SpectraMax M3 microplate reader (BioTek Instruments, Inc., Winooski, VT) at 414 nm within 15 min. The reportable ranges of the E2, T, and P4 assays were 0.61-10,000, 3.9-500, and 7.8-1,000 pg/mL, respectively. The lower limits of detection (LOD) for E2 and T were 6 and 5 pg/mL, respectively (no reported LOD for P4). The sensitivity of hormones, as defined by the manufacturer as the analyte concentration corresponding to 80% B/B_0_ (% Bound/Maximum Bound), and is derived from the standard curve provided in the kit manual, was 20 pg/mL, 10 pg/mL, and 6 pg/mL for E2, P4, and T, respectively. Inter-assay variability (CV) for E2, P4, and T was 8%-12.3%, 7.7%-16.4%, and 7.7%-14.2%, respectively; intra-assay CV for E2, P4, and T was 6.5%-10.8%, 7.3%-54.5%, and 4.4%-19.1%, respectively. Cross-reactivity among E2, P4, and T was less than 0.14%. Experimental controls, blanks, and a series of positive standards were measured every time to ensure the accuracy of hormone levels. N=12-13 follicles were used in each treatment group.

For *in-vivo* hormone analysis, commercial ELISA kits were used (E2: ALPCO; T: IBL) and tested at the University of Virginia Center for Research in Reproduction Ligand Assay and Analysis core. The assay method was validated by spiking steroid-free media with known concentrations of hormone to assess recovery and ensure parallelism with the kit’s standard curve. The assay ranges for T and E2 were 0.15-16.0 ng/mL and 20-3200 pg/mL respectively. The sensitivity of T and E2 were 0.09 ng/ml and 10 pg/ml respectively. The inter-assay CV for T and E2 were 4.7-9.9% and 6.2-10.1% respectively. The intra-assay CV for the same hormones were 3.3-4.2% and 4.6-9.3% respectively. A blank media standard curve was used for each assay to calculate the sample concentrations.

### RNA extraction and RT-qPCR

For *in-vitro* experiments using C-WFPM_0.1_ and LS-WFPM_0.1_, follicles were collected on day 6 of eIVFG and oocytes were removed. Somatic cells from each follicle were used to examine the expression of genes related to follicular growth, steroidogenesis, AhR and NRF2 signaling, and oxidative stress. For *in vivo* animal exposure, follicular somatic cells from three isolated large antral follicles in each mouse ovary (N=3) were pooled together. Total RNA of each follicle or pooled follicular somatic cells was extracted using the PicoPure RNA isolation kit (Thermo Fisher Scientific). RNA purity (A260/A280 ratio) and RNA quantification were performed using NanoDrop. Total RNA was then reversely transcribed into cDNA using the Superscript III First-Strand Synthesis System with random hexamer primers (Invitrogen, catalogue: 18080400) and stored at −80 °C. RT-qPCR was performed in 384-well plates using the Power SYBR Green PCR Master Mix (Thermo Fisher Scientific) in a ViiA 7 Real-Time PCR System (Thermo Fisher Scientific). RT-qPCR thermocycle was programmed for 10 min at 95 °C, followed by 40 cycles of 15 second (s) at 95 °C and 40 s at 60 °C, and ended with a melting curve stage to determine the specificity of primers. The relative gene expression levels of each gene in LS- and C-WFPM_0.1_ exposure experiments were normalized by the expression of glyceraldehyde-3-phosphate dehydrogenase (*Gapdh*), and benzo(a)pyrene (BaP) exposure experiments were normalized by the expression of beta actin (*Actb*). Genes related to steroidogenesis (*Star*, *Cyp11a1*, *Cyp17a1*, *Cyp19a1*, *Hsd3b1*, and *Hsd17b1*), AhR signaling (*Ahr, Ahrr, Cyp1a1, Cyp1b1, Cyp1a2*), oxidative stress and NRF2 signaling (*Nrf2, Sod1, Nqo1*) were examined. For *in vitro* exposure studies, N=5-10 follicles were included per treatment group. The primer sequences of all examined genes are listed in Table S13.

### Single-follicle RNA-sequencing (RNA-seq) analysis

The impact of LS- and C-WFPM_0.1_ on follicular transcriptomic profiles during FSH-induced follicle maturation was evaluated using RNA-seq. Follicles were treated with either vehicle (N=5) or 100 μg/mL of LS-WFPM_0.1_ (N=6) or C-WFPM_0.1_ (N=6) from day 4 to day 6 of eIVFG. At the end of the exposure period, follicles were collected, and total RNA was extracted using the Arcturus PicoPure RNA Isolation Kit (Applied Biosystems), following the manufacturer’s protocol. Library preparation and low-input RNA-seq were conducted on the Illumina NovaSeq PE150 platform (Novogene Corporation, Sacramento, CA). High-quality, trimmed paired-end reads were processed using Partek Flow software. Potential ribosomal DNA (rDNA) and mitochondrial DNA (mtDNA) contaminants were removed using Bowtie 2. The cleaned reads were aligned to the mouse reference genome (mm10) using the HISAT 2 aligner. Raw read counts were generated by mapping aligned reads to Ensembl Transcripts release 99 using the Partek EM algorithm, and normalization was performed using the Transcripts Per Million (TPM) method. Pseudogenes were excluded based on the HUGO Gene Nomenclature Committee’s list of protein-coding genes [102].

Principal component analysis (PCA) was conducted using the PCAtools package in R to assess sample clustering [103]. Differential gene expression analysis was carried out using DESeq2 (R), with significantly differentially expressed genes (DEGs) defined as those with an absolute fold change ≥ 2.0 or ≤ 0.5 and a false discovery rate (FDR)-adjusted *p*-value < 0.05. Kyoto Encyclopedia of Genes and Genomes (KEGG) pathway analysis and Gene Ontology (GO) enrichment analyses were performed on the DEGs using various Bioconductor packages in R [104–106]. Pathway analysis for DEGs obtained from *in-vivo* experiments was performed using WEB-based Gene Set Analysis Toolkit (WebGestalt) [107] and plotted using the ggplot2 package in R [108].

### Single-oocyte SMART-Seq2 RNA sequencing

Oocytes from Day-6 follicles were subjected to single-oocyte Smart-Seq2 RNA sequencing (RNA-seq) using the Smart-seq2 v4 kit (Takara Bio USA Inc.) for library preparation with minor modification from the manufacturer’s instructions. Briefly, single oocytes were lysed, and mRNA was extracted and reverse transcribed into cDNA. cDNA products were amplified by PCR and purified with AMPure XP beads (Beckman Coulter), and further quality-checked by Agilent High Sensitivity DNA assay (Agilent Technologies). High-quality cDNA products were subjected to tagmentation and library preparation, with multiplexing by Nextera XT Indexes (Illumina). Pooled indexed libraries were sequenced on the NovaSeq X Plus platform with 150 bp pair-end reads by Novogene. After sequencing, fastq files were imported into Galaxy (version 24.2.rc1) for downstream analysis. RNA-seq reads were aligned to the whole mouse genome assembly-mm10 by HISAT2, quantified by featureCounts, and normalized using the TPM method. PCA was performed using R (version 4.3.3) and DEG analysis was conducted using DESeq2(R).

### RNA-seq data availability

All the raw FASTQ files and raw counts data from single-follicle and single-oocyte RNA-seq were deposited to the Gene Expression Omnibus (GEO) database with accession numbers of GSE317094, GSE317095, GSE317100 and GSE317185.

### *In situ* RNA hybridization

Follicles collected on day 6 of eIVFG from both control and LS-WFPM_0.1_-treated groups were fixed, paraffin-embedded, and sectioned at a thickness of 5 μm. To assess the spatial expression of the canonical AhR target gene *Cyp1a1*, RNA in situ hybridization was conducted using the RNAscope Multiplex Fluorescent Detection Kit V2 in conjunction with the HybEZ Hybridization System (Advanced Cell Diagnostics, Newark, CA), following the manufacturer’s protocol. Tissue sections underwent a series of pretreatment steps, including heat treatment, hydrogen peroxide incubation, and protease digestion, prior to hybridization with target gene-specific probes. Signal amplification was achieved using an HRP-based system followed by fluorescent dye development. Slides were mounted with Vectashield antifade mounting medium containing DAPI (Maravai LifeSciences, San Diego, CA) and imaged using a Leica confocal microscope (Wetzlar, Germany).

### Data and Statistical Analysis

Statistical analyses were conducted using GraphPad Prism version 10 (GraphPad Software, San Diego, CA). Data distributions were assessed for normality tests and homogeneity of variances before prior to applying the statistical analysis method. For comparisons of numerical data between two treatment groups, such as the gene expression between vehicle-treated and 100 μg/mL real-world or lab-synthesized WFPM-treated follicles, Student’s t-test was applied. One-way ANOVA followed by Dunnett’s multiple comparisons test was used to analyze multiple-group comparisons of numerical data, including follicle diameter and hormone concentrations. Fisher’s exact test was utilized to assess categorical data, such as *in vitro* follicle rupture and oocyte polar body extrusion. The hormone concentration data was log-transformed to normalize data distributions and meet assumptions of parametric analysis prior to ANOVA testing. Data are presented as mean ± standard deviation (SD), and statistical significance was defined as a *p*-value less than 0.05. For *in vitro* hormone dose-response data that were statistically significant, Bayesian benchmark dose (BBMD) analysis was applied to derive benchmark concentrations (BMC) by using the default parameter setting and model averaging for continuous data [109]. The BBMD is a more robust platform than the US EPA Benchmark Dose Modeling Software (BMDS) tool [110]. A relative deviation of 10% from the background level was set as the benchmark response (BMR_10_) to calculate the median benchmark concentrations (BMC_10_) and corresponding 95% lower confidence limit (BMCL_10_) of the WFPM solutions.

## Results

### Physicochemical characterization of LS- and C-WFPM_0.1_

Physicochemical characterization of LS-WFPM_0.1_ has been reported in detail in our companion paper from Laurent et al. [8]. In brief, organic carbon and elemental carbon analysis (OC/EC) showed that there was a dominance of organic carbon in both LS- and C-WFPM_0.1_, with the organic to total carbon (OC/TC) ratios of ∼ 99.7% and 96%, respectively (Figure S2A and S2D). These high organic-to-inorganic ratios are consistent with values reported for biomass combustion [111, 112] and confirm the organic-rich composition of WFPM_0.1_.

Elemental analysis revealed that both LS- and C-WFPM_0.1_ were dominated by crustal elements, with iron (Fe) representing the most abundant component in both particle types (Figure S2B and S2E). Aluminum (Al), titanium (Ti), and manganese (Mn) were also detected in both samples, indicating shared elemental features characteristic of mineral-rich WFPM_0.1_. LS-WFPM_0.1_ also contained alkaline earth metals, with low levels of trace metals. This elemental composition is consistent with a controlled combustion environment and the absence of soil or anthropogenic contributions [8]. C-WFPM_0.1_ exhibited a broader array of trace elements, including barium (Ba), lead (Pb), manganese (Mn), zirconium (Zr), and zinc (Zn), which are often associated with crustal sources and anthropogenic sources such as traffic and industrial emissions, reflecting the environmental mixing and long-range transport characteristics of the real-world wildfire plumes.

LS- and C-WFPM_0.1_ exhibited both similarities and differences in their PAH profiles (Figures S2F and S2C). Retene, a marker of wood combustion [35], was the predominant PAH in both LS- (∼97.91%) and C-WFPM_0.1_ (∼76.73%). Both types of WFPM_0.1_ contained other medium (4 aromatic rings) and high (> 4 aromatic rings) molecular weight PAHs, including benzo(e)pyrene, fluoranthene, pyrene, chrysene, indeno(1,2,3-cd)pyrene, and benzo(g,h,i)perylene. LS- and C-WFPM_0.1_ also exhibited distinctions in their PAH profiles, with benzo(a)pyrene (BaP) present only in LS-WFPM_0.1_. Further characterization results of WFPM_0.1_ *in-vitro* colloidal properties, as well as endotoxin and sterility analysis, are given in Supplementary Information. Collectively, these findings indicate that although there are distinctions between LS- and C-WFPM_0.1_, primarily due to other sources in the real-world PM samples, LS-WFPM_0.1_ reproduces key elemental fingerprints of the real-world WFPM_0.1_.

### Intratracheal exposure to LS-WFPM_0.1_ interferes with mouse ovarian cycles and steroidogenesis

From day 1 to 7 post-instillation, mice treated with vehicle or LS-WFPM_0.1_ had comparable survival, behavior, body weight, and water and food consumption (results not shown). Biochemical analysis of BALF showed that on day 7 post-exposure, mice treated with LS-WFPM_0.1_ had ∼1.9-fold higher concentrations of lactate dehydrogenase (LDH), a tissue damage marker, compared to control (Figure S3), indicating lung cell injury caused by LS-WFPM_0.1_, consistent with prior studies [31, 32, 113–115].

Daily vaginal smears revealed that mice exposed to LS-WFPM_0.1_ spent significantly fewer days on proestrus (*p*=0.04) and a borderline longer duration on diestrus (*p*=0.09) (Figure 1B). Mice treated with LS-WFPM_0.1_ had slightly higher serum concentrations of both estradiol and testosterone, but the differences were not statistically significant (Figure 1C), even focusing on proestrus alone (Figure 1D). Antral follicles were isolated from mice treated with vehicle or LS-WFPM_0.1_ on day 7 to examine steroidogenic gene expression by RT-qPCR. Mice treated with LS-WFPM_0.1_ tended to have higher expressions of *Star*, *Cyp11a1*, and *Hsd3b1* and lower expressions of *Cyp17a1* and *Hsd17b1,* although differences were not statistically significant (Figure 1E). Expression of *Cyp19a1*, encoding aromatase, was comparable. Collectively, these results suggest that intratracheal exposure to LS-WFPM_0.1_ interferes with mouse ovarian cycles and steroidogenesis.

### Intratracheal exposure to LS-WFPM_0.1_ activates AhR in ovarian antral follicles

Our recent WFPM exposure assessment studies revealed PAHs as a major organic component of WFPM_0.1_ [14, 74]. PAHs have been reported to act as AhR agonists to exhibit endocrine disrupting effects [116–118]. We thus examined several AhR target genes in antral follicles using RT-qPCR, including *Ahr*, *Cyp1a1*, *Cyp1b1*, *Ahrr*, *Nrf2*, and *Nqo1* [119]. The full name, major functions, and related references are summarized in Table S1. LS-WFPM_0.1_ did not affect the expression of *Cyp1a1*, *Cyp1b1*, and *Ahrr*, but significantly increased *Ahr* and *Nrf2* by 1.50- and 1.27-fold, respectively. *Nqo1* exhibited a slight increase (Figure 1F). These results suggest that LS-WFPM_0.1_ activates the AhR signaling pathway in antral follicles.

### Intratracheal exposure to LS-WFPM_0.1_ alters follicular somatic cell transcriptome

To determine the effect of LS-WFPM_0.1_ on the follicular transcriptome, we used total RNA isolated from antral follicular somatic cells for RNA-seq analysis. PCA moderately separated antral follicles from vehicle or LS-WFPM_0.1_-treated mice (Figure 2A). There were 76 DEGs with fold change >= 2 or <= 0.5 and FDR < 0.05, including 48 up- and 28 down-regulated genes in LS-WFPM_0.1_-treated follicles, with the top 10 up-and down-regulated DEGs highlighted in the volcano plot and heatmap in Figure 2A. All DEGs are listed in Excel Table S2. The most upregulated gene was *Ces2a*, which encodes a carboxylesterase enzyme that regulates lipid, drug, and xenobiotic metabolism [120, 121]. The most downregulated gene was *Fosb*, which encodes the Fos protein that heterodimerizes with Jun to regulate important cellular processes such as proliferation and differentiation [122]. Similar to RT-qPCR data, RNA-seq data showed consistent changes in most ovarian steroidogenic genes and AhR target genes (Figure 2B), including increases in *Star, Cyp11a1, Hsd3b1, Ahr, Nrf2,* and *Nqo1*. *Ahrr*, another established AhR target gene was slightly increased in LS-WFPM_0.1_-treated follicles.

**Figure 2.**
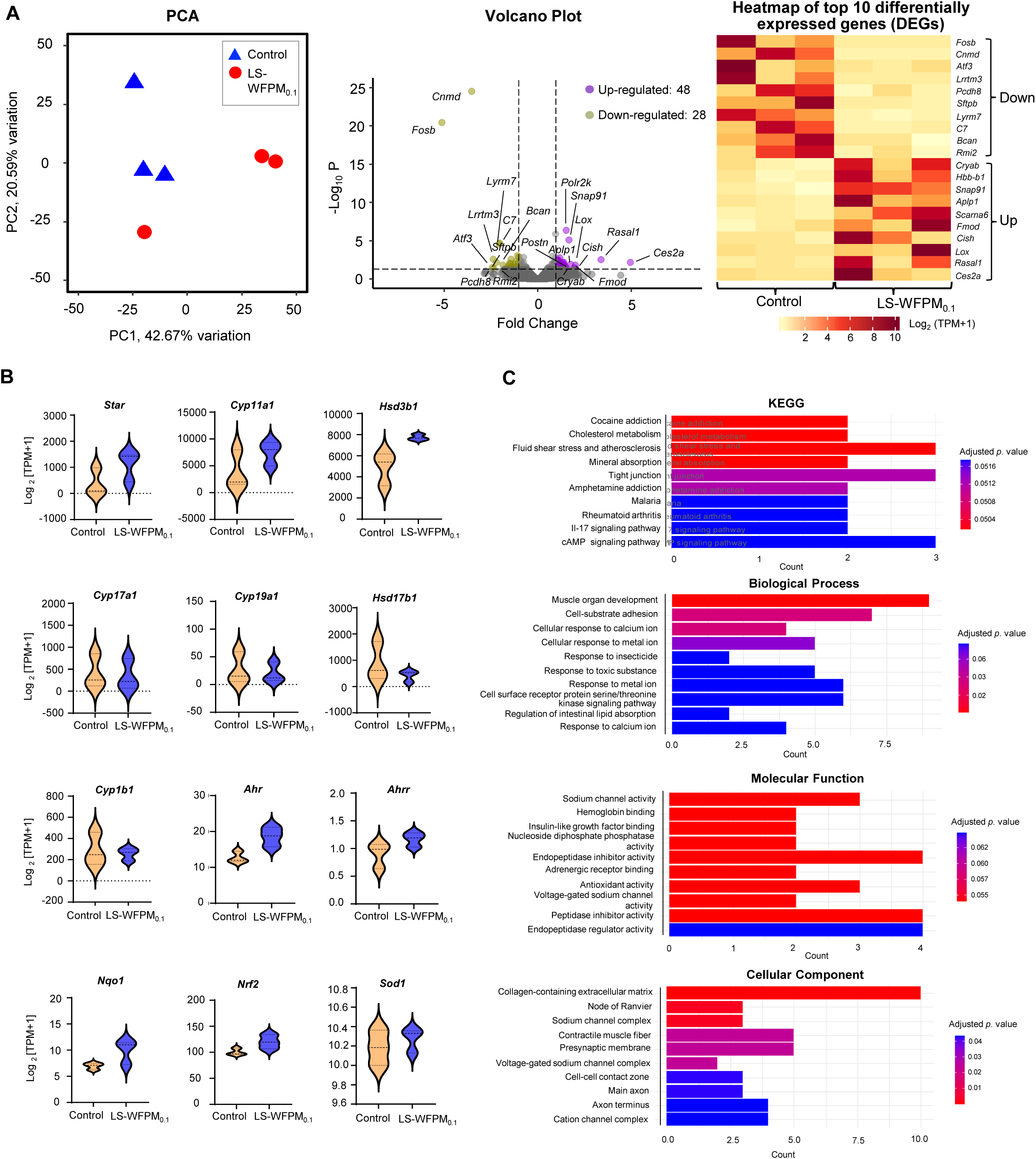
RNA-seq analysis of 3 pooled antral follicles derived from control (N=3) or LS-WFPM_0.1_ treated (N=3) mice collected on day 7 post-exposure. (A) PCA of the first two PCs for control or LS-WFPM_0.1_ follicles; Volcano plot of DEGs (labeled genes have FDR < 0.05, absolute fold change >= 2 or <=0.5) in LS-WFPM_0.1_ treated follicles compared to the control; Heatmap of the top 10 up- and down-regulated genes as indicated. Each column in the heatmap represents the relative difference of gene expression level in each treatment group. (B) Violin plots of mRNA expression levels in Log_2_ [Transcript Per Million (TPM) +1] showing major steroidogenesis, AhR-related and oxidative stress-related genes in control and LS-WFPM_0.1_ antral follicles. (C) KEGG pathway and GO analysis of DEGs identified by RNA-seq between 3 pooled follicles derived from control or LS-WFPM_0.1_ treated mice, including top 10 enriched KEGG pathways, biological processes, molecular functions and cellular component as indicated. X-axes in KEGG pathway and GO analysis represent the number of genes that were present in the list of DEGs and associated with that pathway or GO term. Data presented in (C) are also provided in Excel Table S3 and S4 for GO terms and KEGG pathways respectively.

DEGs were used for GO and KEGG pathway analyses. All enriched GO terms and pathways are listed in Excel Tables S3 and S4, and the top 10 are highlighted in Figure 2C. Biological process analysis revealed DEGs associated with “Muscle organ development”, “Cell-substrate adhesion”, and “Cellular response to calcium ion” as the top altered processes. Molecular function analysis showed DEGs involved in “Sodium channel activity”, “Hemoglobin binding”, and “Insulin-like growth factor binding”. Cellular component analysis discovered DEGs associated with “Collagen-containing extracellular matrix”, “Node of Ranvier”, and “Sodium channel complex”. KEGG analysis revealed that DEGs are related to “Cocaine addiction”, “Cholesterol metabolism”, and “Mineral absorption”.

### LS-WFPM_0.1_ induces ovarian hyperandrogenism *in vitro*

To elucidate the mechanisms of perturbed folliculogenesis and steroidogenesis caused by LS-WFPM0.1, we used a 3D hydrogel eIVFG model for in vitro exposure studies. Immature mouse secondary follicles were treated with 0, 1, 10, and 100 µg/mL LS-WFPM0.1 from day 4 to 6 in eIVFG (Figure. 3A). Follicles treated with all concentrations of LS-WFPM0.1 exhibited comparable morphology and survival to the control (Figure. 3B). Follicles from all groups were able to grow from the secondary to antral stage with comparable follicle sizes (Figure. 3C). Upon ovulation induction in vitro on day 6, most follicles from vehicle and LS-WFPM0.1 groups ruptured and ovulated MII oocytes with the first polar body extrusion on day 7 (Figure 3C). Follicles from both groups had comparable estradiol secretion on day 6 and progesterone secretion on day 9 (Figure 3D). However, consistent with our in vivo findings, LS-WFPM0.1 increased testosterone secretion in a concentration-dependent manner, reaching 1.3-fold at 100 µg/mL that is statistically significant (Figure 3D).

**Figure 3.**
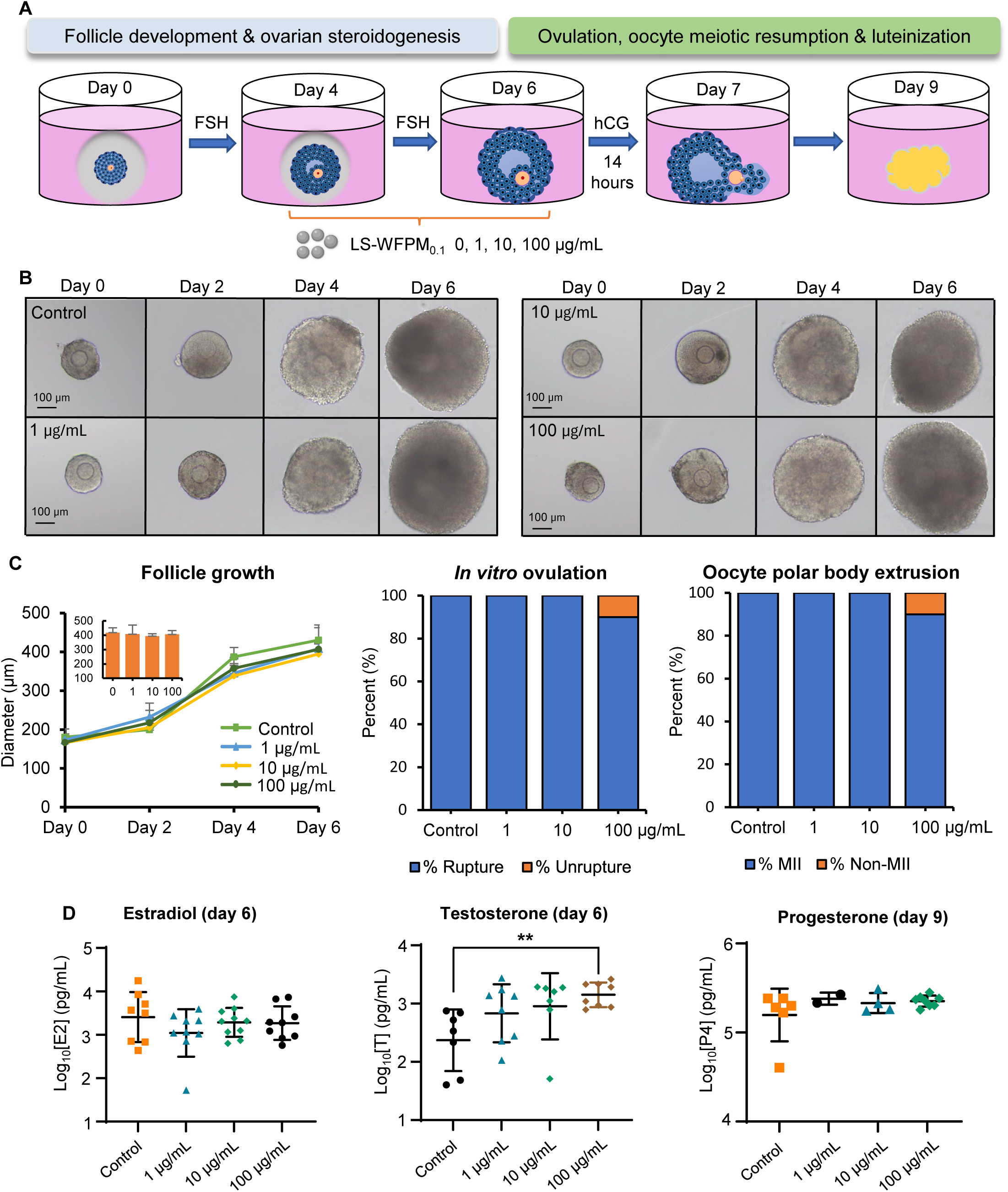
Effects of *in vitro* exposure to LS-WFPM_0.1_ during gonadotropin dependent follicle maturation window (day 4-6) on follicle growth, ovulation and hormone secretion. (A) Schematic of LS-WFPM_0.1_ exposure using 3D eIVFG model. (B) Representative follicle images from day 0 to day 6 of eIVFG from control and LS-WFPM_0.1_ exposed group with the indicated doses. (C) Follicle diameter of control and LS-WFPM_0.1_ exposed group (N=10-12 follicles per group) at various concentrations; Percent of ruptured, un-ruptured follicles and ovulated MII oocytes on day 7 of eIVFG. (D) Average Log_10_ concentration (pg/mL) of estradiol and testosterone in the conditioned follicle culture media collected on day 6 of eIVFG (N=7-10 follicles per group); Progesterone concentrations (pg/mL) in follicle culture media collected on day 9 of eIVFG (N=2-8 follicles per group). Data were analyzed with one-way ANOVA followed by Dunnett’s multiple comparisons test. Error bars represent mean ± standard deviation. \*\**p < 0.01.* For *in vitro* ovulation and polar body extrusion in (C), data were analyzed using Fischer’s exact test.

### LS-WFPM_0.1_ alters genes related to follicle maturation, steroidogenesis, and AhR signaling *in vitro*

Follicles treated with vehicle or 100 µg/mL LS-WFPM_0.1_ were collected on day 6 to examine the panels of genes related to steroidogenesis and AhR using RT-qPCR. Consistent with *in vivo* data, LS-WFPM_0.1_ exhibited similar effects on steroidogenic genes, including decreased expressions of *Cyp17a1* and *Hsd17b1*, along with an upward trend in *Hsd3b1* expression. Different from *in vivo* under which *Cyp19a1* expression remained unchanged, follicles treated with LS-WFPM_0.1_ *in vitro* exhibited a borderline decrease in expression of *Cyp19a1* (Figure 4A). Upstream steroidogenic genes, such as *Star* and *Cyp11a1* that were slightly upregulated *in vivo*, remained unchanged (Figure 4A). Most AhR target genes exhibited consistent expression patterns between *in vitro* and *in vivo*, including *Cyp1b1*, *Ahrr*, *Nqo1*, and *Nrf2* (Figure 4B). *Cyp1a1*, an established AhR target gene, was not altered *in vivo* but was markedly induced *in vitro* (Figure 4B). *In situ* RNA hybridization was used to examine *Cyp1a1* using follicles treated with vehicle or 100 µg/mL LS-WFPM_0.1_. *Cyp19a1* was used as a marker gene for granulosa cells. Consistent with RT-qPCR data, LS-WFPM_0.1_ significantly increased *Cyp1a1* in both *Cyp19a1*-postive and negative cells in the outer follicle layer, suggesting the activation of AhR in both mural granulosa and theca cells (Figure 4C). Collectively, these results confirm the perturbation of ovarian steroidogenesis and AhR activation in follicles treated with LS-WFPM_0.1_.

**Figure 4.**
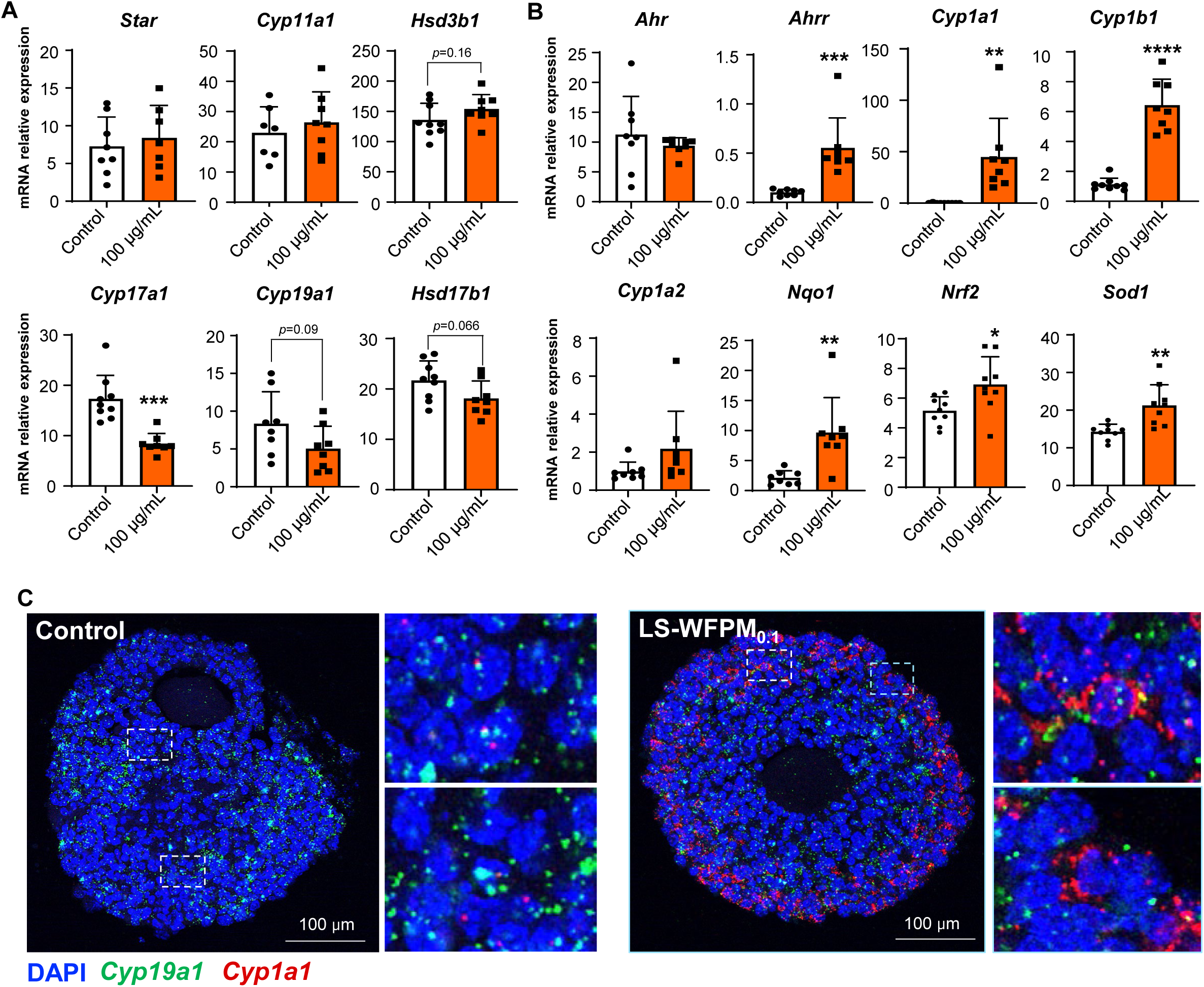
*In vitro* exposure to 100 µg/mL LS-WFPM_0.1_ alters genes related to follicle maturation, steroidogenesis, and AhR signaling. (A) Relative mRNA expression of follicle maturation and steroidogenesis related genes examined by RT-qPCR in control and LS-WFPM_0.1_ treated follicles. (B) Relative mRNA expression of AhR related genes examined by RT-qPCR in control and LS-WFPM_0.1_ treated follicles. N=7-9 follicles per group. Expression levels were normalized by the expression of *Gapdh*. Data were analyzed with Student’s t-test. Error bars represent mean ± standard deviation. \**p < 0.05*, \*\**p < 0.01,* \*\*\**p < 0.001.* (C) Representative images of *in situ* hybridization of large antral follicles for *Cyp19a1* and *Cyp1a1* in control and LS-WFPM_0.1_ group.

### LS-WFPM_0.1_ alters the follicular transcriptome and activates AhR signaling

To identify gene regulatory pathways responsible for the ovarian disrupting effects of LS-WFPM_0.1_, follicles treated with vehicle or 100 µg/mL LS-WFPM_0.1_ were collected on day 6 of eIVFG. Follicular somatic cells and oocytes were separated for single-follicle RNA-seq analysis and single-oocyte SMART-seq2 RNA-seq analysis, respectively. The RNA-seq data of follicular somatic cells showed that PCA separated follicles treated with vehicle and LS-WFPM_0.1_ into two distinct clusters (Figure 5A), indicating a significantly altered follicular transcriptome. There were 5,427 DEGs with fold change >= 2 or <= 0.5 and FDR < 0.05, including 2,646 up- and 2,781 down-regulated genes induced by LS-WFPM_0.1_ (Figure 5A). All DEGs are listed in Excel Table S5. The top 10 up- and down-regulated genes are highlighted in the volcano plot and heatmap in Figure 5A. The most upregulated gene was the classical AhR target *Cyp1a1*, showing a 2,465-fold increase, while another AhR-responsive gene, *Cyp1b1*, was also among the top ten upregulated genes with an 11-fold increase, together confirming strong activation of AhR signaling in follicles treated with LS-WFPM_0.1_. Conversely, the most downregulated gene was *Tnn*, encoding tenascin-N/tenascin-W, an extracellular matrix glycoprotein with unknown ovarian functions [123]. Consistent with the RT-qPCR data (Figure 4A, 4B), the same set of genes exhibited similar changes in the RNA-seq analysis (Figure 5B). Other AhR target genes, including *Ahrr*, *Nrf2*, and *Nqo1*, were also significantly increased in follicles treated with LS-WFPM_0.1_ (Figure 5B). Several oxidative stress and NRF2-related genes, including *Sod1*, *Sod2*, *Gsr*, *Gstp1*, and *Gpx4*, were significantly increased in LS-WFPM_0.1_-treated follicles.

**Figure 5.**
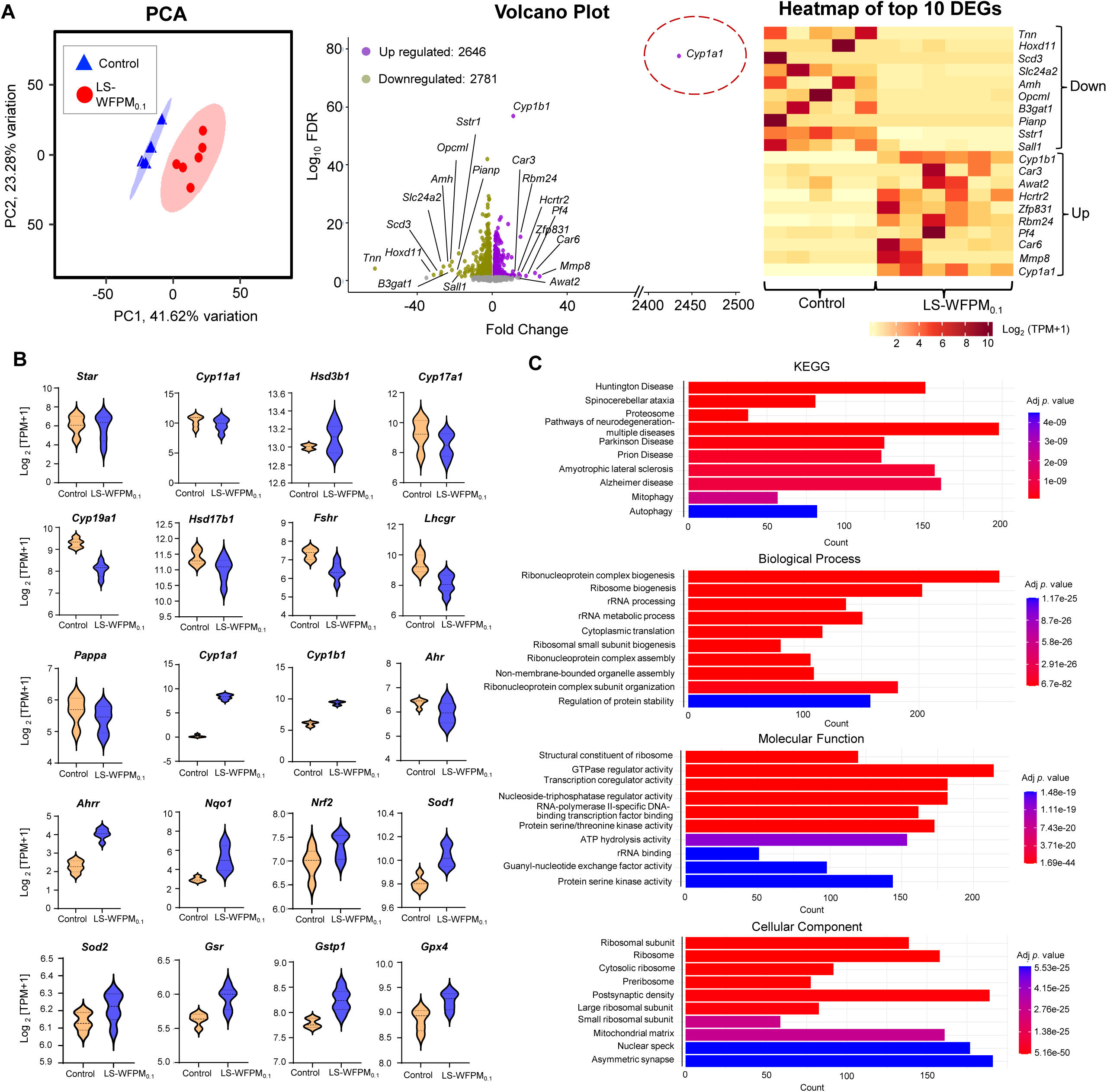
RNA-seq analysis of single antral follicles derived from control (N=5) or 100 µg/mL LS-WFPM_0.1_ treatment (N=6) collected on day 6 of eIVFG. (A) PCA of the first two PCs for control or LS-WFPM_0.1_ follicles; Volcano plot of DEGs (top 10 gene with FDR < 0.05 and absolute fold change >= 2 or <=0.5 are labeled) in LS-WFPM_0.1_ treated follicles compared to the control; Heatmap of the top 10 up- and down-regulated genes as indicated. Each column in the heatmap represents the relative difference of gene expression level in each treatment group. (B) Violin plots of mRNA expression levels in Log_2_ [TPM+1] showing major steroidogenesis, follicle maturation, AhR-related and oxidative stress related genes in control and LS-WFPM_0.1_ antral follicles. (C) KEGG pathway and GO analysis of DEGs identified by single-follicle RNA-seq between control or LS-WFPM_0.1_ treated follicles during the follicle maturation window, including top 10 enriched KEGG pathways, biological processes, molecular functions and cellular component as indicated. X-axes in KEGG pathway and GO analysis represent the number of genes that were present in the list of DEGs and associated with that pathway or GO term. Data presented in (C) are also provided in Excel Table S6 and S7 for GO terms and KEGG pathways respectively.

DEGs were used for GO term enrichment and KEGG pathway analyses. All significantly enriched GO terms and pathways are listed in Excel Tables S6 and S7, respectively, and the top 10 are highlighted in Figure 5C. Biological process analysis showed that DEGs were primarily related to “Ribonucleoprotein complex biogenesis”, “Ribosome biogenesis”, and “rRNA processing” (Figure 5C). Molecular function analysis enriched DEGs associated with “Structural constituent of ribosome”, “Transcription coregulator activity”, and “GTPase regulator activity” (Figure 5C). Cellular component analysis showed DEGs involved in “Ribosomal subunit”, “Ribosome”, and “Cytosolic ribosome” (Figure 5C). KEGG analysis revealed DEGs related to “Huntington disease”, “Spinocerebellar ataxia”, and “Proteosome” (Figure 5C).

### LS-WFPM_0.1_ alters the oocyte transcriptome and genes related to developmental competence without activating AhR signaling

Oocytes were collected on day 6 of eIVFG for single-oocyte SMART-Seq2 RNA-seq analysis to determine the effect of LS-WFPM_0.1_ on the quality of follicle-enclosed oocytes. PCA moderately separated oocytes from vehicle and LS-WFPM_0.1_-treated follicles (Figure 6A), suggesting a change in the oocyte transcriptome. The expression of key oocyte-specific genes associated with oocyte growth and identity, including *Gdf9*, *Bmp15*, and zona pellucida genes (*Zp1*, *Zp2*, and *Zp3*), was comparable between control and LS- WFPM_0.1_-exposed follicles (Figure 6B, left panel). Different from follicular somatic cells, there were no AhR target genes elevated in oocytes from LS-WFPM_0.1_-treated follicles (Figure 6B, right panel). There were 846 DEGs, including 434 up- and 412 down-regulated genes in oocytes from LS-WFPM_0.1_-treated follicles. The GO term and KEGG analyses further identified DEGs associated with “oxidative phosphorylation”, “mitochondrial respiratory chain”, “Ubiquitin protein ligase binding”, “protein catabolic process”, and “NADH dehydrogenase complex” (Figure 6C). When comparing all DEGs to 82 mammalian maternal effect genes (MEGs) [124], there were 5 genes that overlapped, including 3 up-regulated MEGs (*Zbed3*, *Mapk3*, and *Khdc3*) and 2 down-regulated MEGs (*Dappa3* and *Ago2*). The gene names, fold-change, and major biological functions are summarized in Table S8.

**Figure 6.**
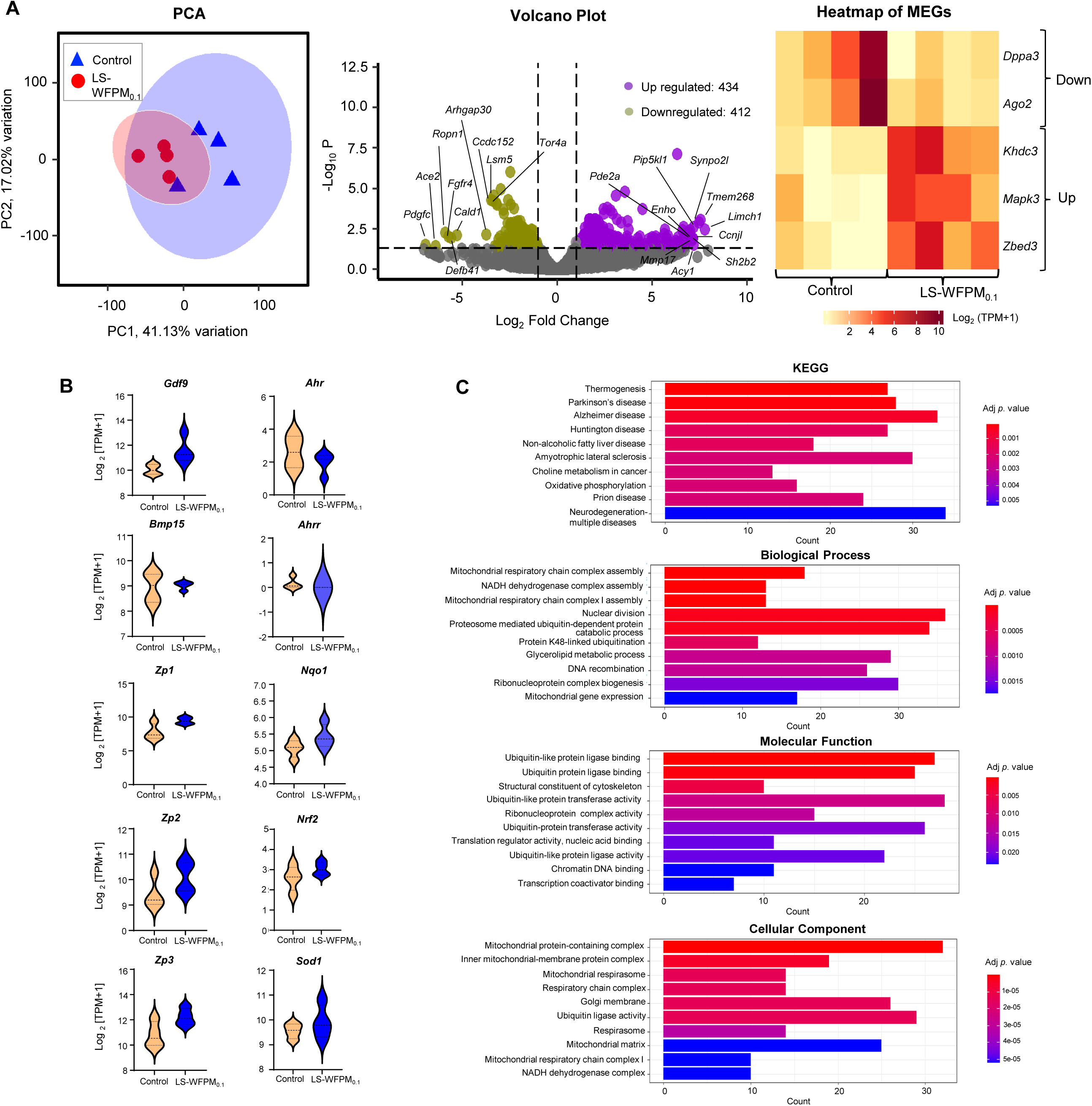
SMART-Seq2 RNA-seq analysis of single oocyte derived from antral follicles in control (N=4) or 100 µg/mL LS-WFPM_0.1_ treated group (N=4) collected on day 6 of eIVFG. (A) PCA of the first two PCs for control or LS-WFPM_0.1_ follicle derived oocyte; Volcano plot of DEGs (top 10 genes with FDR < 0.05 and absolute fold change >= 2 or <=0.5 are labeled) in LS-WFPM_0.1_ treated oocyte compared to the control; Heatmap of the top 10 DEGs. (B) Violin plots of mRNA expression levels in Log_2_ [TPM+1] showing major oocyte marker genes, AhR-related and oxidative stress-related genes in control and LS-WFPM_0.1_ follicle derived oocytes (C) KEGG pathway and GO analysis of DEGs identified by SMART-Seq2 RNA-seq between control or LS-WFPM_0.1_ treated follicle derived oocytes during the follicle maturation window, including top 10 KEGG pathways, biological processes, molecular functions and cellular component as indicated.

### C-WFPM_0.1_ exhibits similar ovarian disrupting effects to LS-WFPM_0.1_

We next used eIVFG to test the real-world WFPM_0.1_ collected from the central New Jersey area during the Canadian wildfire event in June 2023. The exposure strategy was similar to the LS-WFPM_0.1_ exposure in Figure 3A. C-WFPM_0.1_ did not affect follicle survival (Figure 7A), growth, ovulation (Figure 7B), and estradiol secretion (Figure 7C), but it consistently resulted in ovarian hyperandrogenism. Unlike LS-WFPM_0.1,_ which did not alter progesterone, C-WFPM_0.1_ inhibited progesterone secretion at 100 µg/mL in the formed corpus luteum (CL) organoids on day 9 (Figure 7C). RT-qPCR using cultured follicles from day 6 showed that several steroidogenic genes were consistently downregulated, including *Cyp19a1* and *Hsd17b1* (Figure 8A). Unlike LS-WFPM_0.1_-treated follicles, where *Cyp11a1* exhibited no change, and *Cyp17a1* was reduced, C-WFPM_0.1_ reduced the expression of *Cyp11a1* and did not alter *Cyp17a1*. *Star* was unchanged, which is consistent with LS-WFPM_0.1_. C-WFPM_0.1_ consistently upregulated most AhR target genes, like *Cyp1a1*, *Cyp1b1*, *Ahrr*, *Nqo1*, and *Nrf2,* which were also significantly upregulated (Figure 8B).

**Figure 7.**
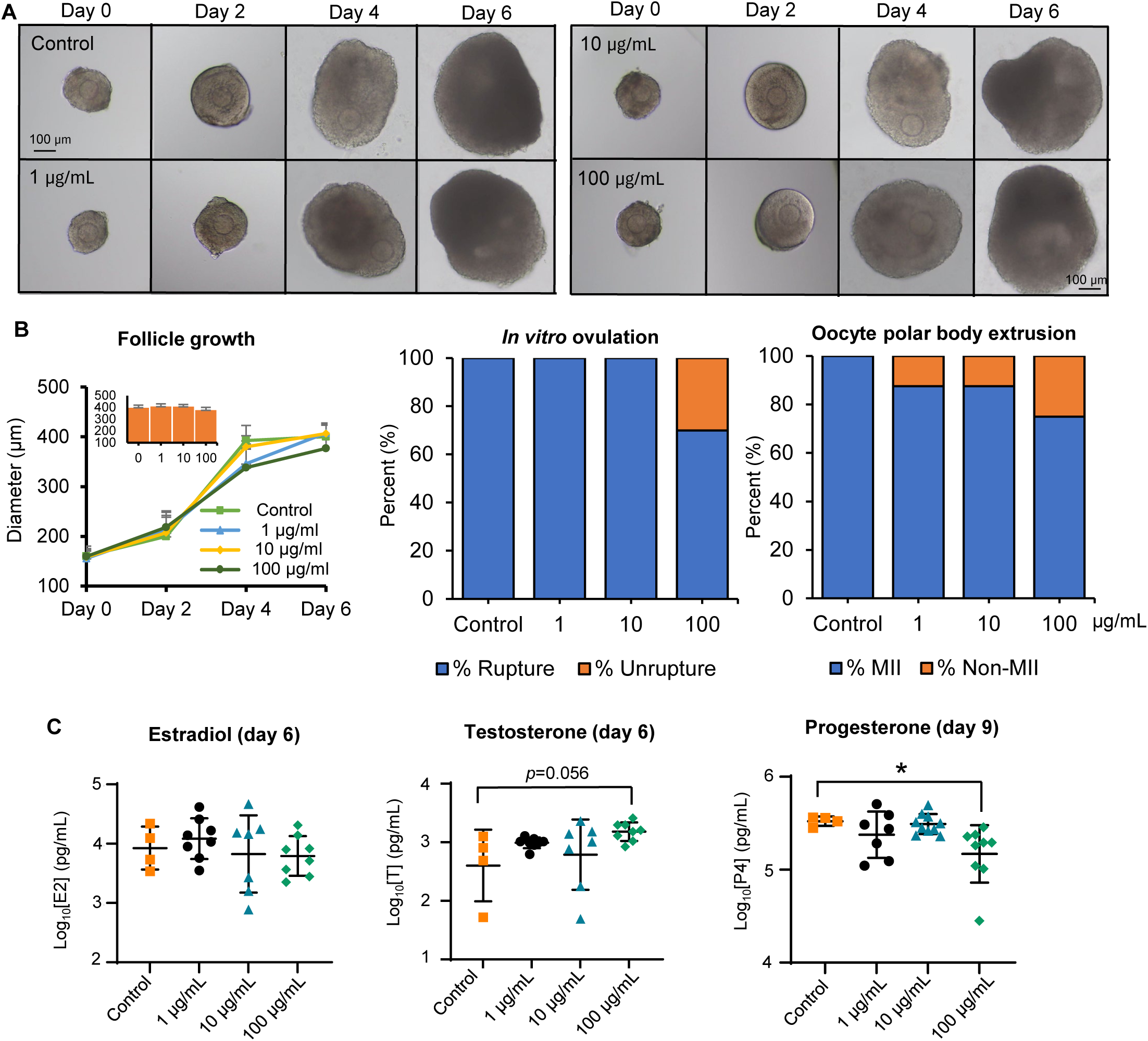
Effects of *in vitro* exposure to C-WFPM_0.1_ during gonadotropin dependent follicle maturation window (day 4-6) on follicle growth, ovulation and hormone secretion. (A) Representative follicle images from day 0 to day 6 of eIVFG from control and C-WFPM_0.1_ exposed group with the indicated doses. (B) Follicle diameter of control and C-WFPM_0.1_ exposed group (N=10-12 follicles per group) at various concentrations; Percent of ruptured, un-ruptured follicles and ovulated MII oocytes on day 7 of eIVFG. (C) Average Log_10_ concentration (pg/mL) of estradiol and testosterone in the conditioned follicle culture media collected on day 6 of eIVFG (N=4-8 follicles per group); Progesterone concentrations (pg/mL) in follicle culture media collected on day 9 of eIVFG (N=4-10 follicles per group). Data were analyzed with one-way ANOVA followed by Dunnett’s multiple comparisons test. Error bars represent mean ± standard deviation. \**p < 0.05.* For *in vitro* ovulation and polar body extrusion in (B), data were analyzed using Fischer’s exact test.

**Figure 8.**
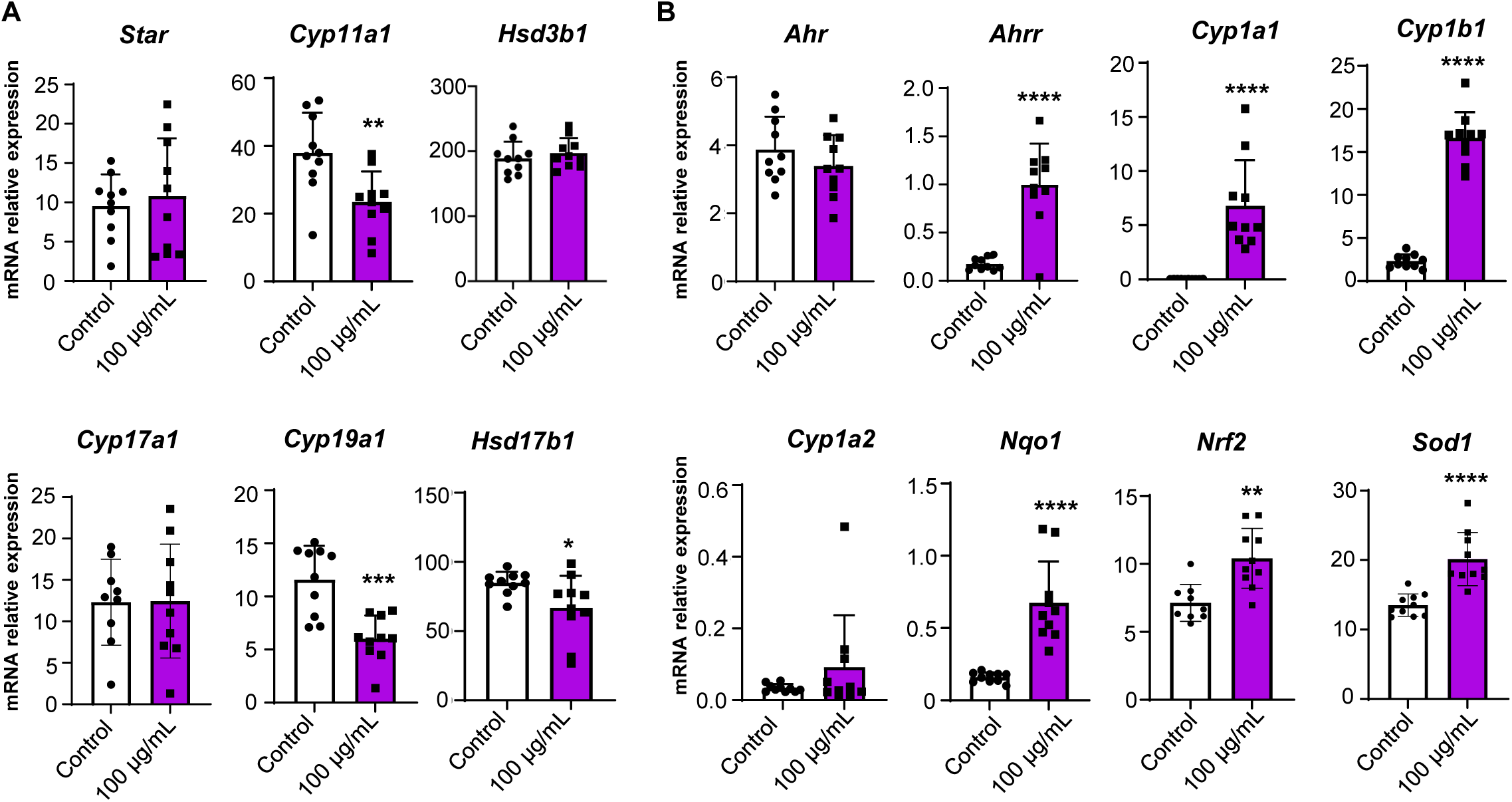
*In vitro* exposure to 100 µg/mL C-WFPM_0.1_ altered genes related to follicle maturation, steroidogenesis, and AhR signaling. (A) Relative mRNA expression of the panels of follicle maturation and steroidogenesis related genes examined by RT-qPCR. (B) Relative mRNA expression of the panels of AhR related genes examined by RT-qPCR. N=10 follicles per group. Expression levels were normalized by the expression of *Gapdh*. Data were analyzed with Student’s t-test. Error bars represent mean ± standard deviation. \**p < 0.05*, \*\**p < 0.01,* \*\*\*\**p < 0.0001*.

Follicles treated with vehicle or 100 µg/mL C-WFPM_0.1_ were collected from day 6 for RNA-seq analysis. PCA separated two groups of follicles (Figure 9A). There were 4,335 DEGs in follicles treated with C-WFPM_0.1_, including 2,338 up- and 1,997 down-regulated genes (Figure 9A). RNA-seq confirmed transcriptional changes of genes examined by RT-qPCR (Figure 9B). DEGs were used for GO term and KEGG pathway analyses. All significantly enriched GO terms and pathways are listed in Excel Tables S9 and S10, respectively, and the top 10 are highlighted in Figure 9C. Biological process analysis identified DEGs related to “Small GTPase mediated signal transduction, “Cell-substrate adhesion”, and “Actin-filament organization”. Molecular function analysis revealed DEGs involved in “GTPase regulator activity”, “Nucleoside-triphosphatase regulator activity”, and “Actin binding”. Cellular component analysis showed that DEGs were related to “Cell leading edge”, “Collagen-containing extracellular matrix”, and “Ruffle”. KEGG analysis identified “AGE-RAGE signaling pathway in diabetic complications” as the most significantly altered signaling in LS-WFPM_0.1_-treated follicles, followed by “Axon guidance” and “Human papillomavirus infection”.

**Figure 9.**
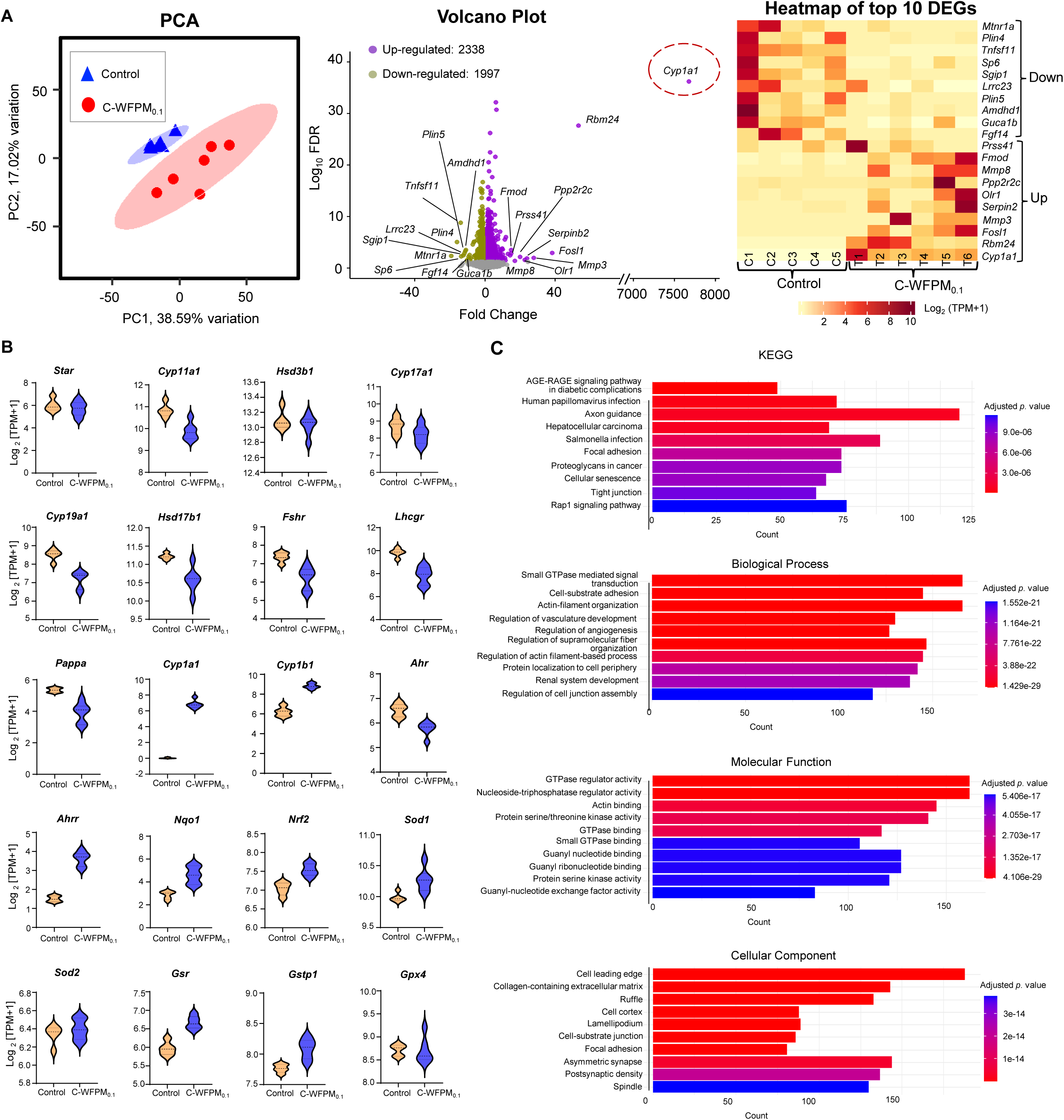
RNA-seq analysis of single antral follicle derived from control (N=5) or 100 µg/mL C-WFPM_0.1_ treatment (N=6) collected on day 6 of eIVFG. (A) PCA of the first two PCs for control or C-WFPM_0.1_ follicles; Volcano plot of DEGs (top 10 genes with FDR < 0.05 and absolute fold change >= 2 or <=0.5 are labeled) in C-WFPM_0.1_ treated follicles compared to the control; Heatmap of the top 10 DEGs examined by RNA-seq analysis. (B) Violin plots of mRNA expression levels in Log_2_ [TPM+1] showing major steroidogenesis, follicle maturation, AhR-related and oxidative stress related genes in control and C-WFPM_0.1_ antral follicles. (C) KEGG pathway and GO analysis of DEGs identified by single-follicle RNA-seq between control or C-WFPM_0.1_ treated follicles during the follicle maturation window, including top 10 enriched KEGG pathways, biological processes, molecular functions and cellular component as indicated. Data presented in (C) are also provided in Excel Table S9 and S10 for GO terms and KEGG pathways respectively.

The transcriptome changes between follicles treated with C- and LS-WFPM_0.1_ were compared. There were 4,335 and 5,427 DEGs, respectively, altered by C-WFPM_0.1_ and LS-WFPM_0.1_, with 1,974 overlapped DEGs, accounting for 45.5% of all DEGs identified in the C-WFPM_0.1_-treated follicles and 36.4% in LS-WFPM_0.1_-treated follicles (Figure 10A). 2,361 DEGs were selectively altered by C-WFPM_0.1_, and 3,453 DEGs by LS-WFPM_0.1_. Correlation analysis calculated a Pearson correlation coefficient value (R) of 0.6 (Figure 10B) of DEGs between follicles treated with the C- and LS-WFPM_0.1._, suggesting a moderate positive correlation as well as distinct transcriptomic changes between the two treatments which could be explained given the differences in their physicochemical properties and potential synergistic, antagonistic effects due to the presence of other chemical compounds (i.e., heavy metals) from other ambient sources as shown above.

**Figure 10.**
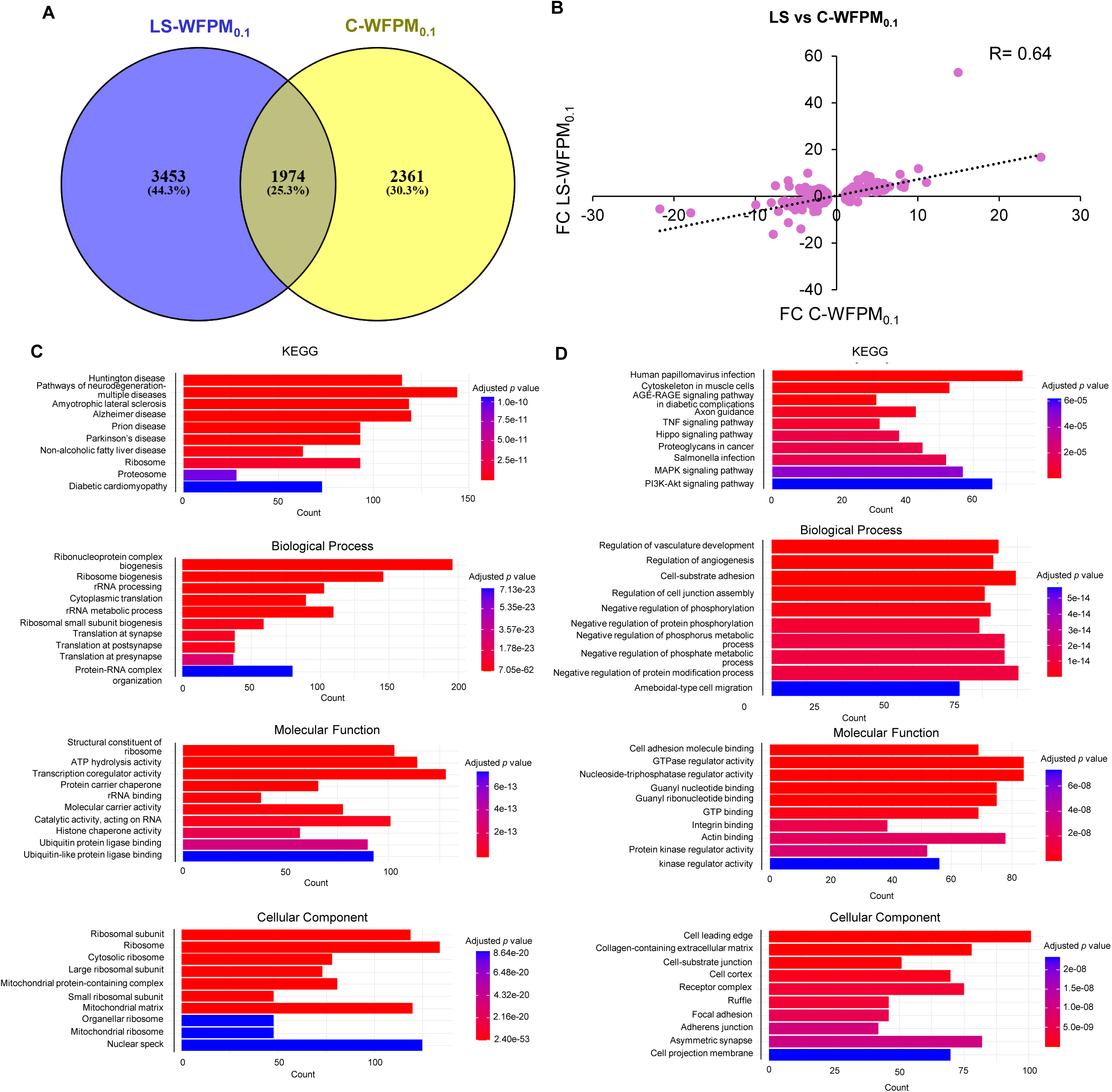
Comparison of 100 µg/mL LS (N=6) and C-WFPM_0.1_ (N=6) treated single follicle RNA-seq analysis. (A) A Venn diagram identifying the number of DEGs overlapping between the treatments of LS-WFPM_0.1_ and C-WFPM_0.1_ as well as those selectively altered by the two treatment types. (B) Pearson correlation analysis of DEGs for LS- and C-WFPM_0.1_ treated single follicles. (C) KEGG pathways and GO analysis of DEGs selectively altered by LS- WFPM_0.1_ (D) KEGG pathways and GO analysis of DEGs selectively altered by C- WFPM_0.1_.

The GO term and KEGG pathway analysis showed that overlapped DEGs were related to “Hepatocellular carcinoma”, “Spinocerebellar ataxia”, “small GTPase-mediated signal transduction”, “Actin filament organization”, “Transcription coregulator activity”, “GTPase regulator activity”, “Cell leading edge”, and “Cell cortex” (Figure S4). DEGs selectively altered by LS-WFPM_0.1_ are associated with “Huntington disease”, “Pathways of neurodegeneration-multiple diseases”, “Ribonucleoprotein complex biogenesis”, “Ribosome biogenesis”, “Structural constituent of ribosome”, “ATP hydrolysis activity”, “Ribosomal subunit”, and “Ribosome” (Figure 10C). DEGs selectively altered by C-WFPM_0.1_ were related to “Human papillomavirus infection”, “Cytoskeleton in muscle cells”, “Regulation of vasculature development”, “Regulation of angiogenesis”, “Cell adhesion molecule binding”, “GTPase regulator activity”, “Cell leading edge”, and “Collagen-containing extracellular matrix” (Figure 10D). These bioinformatic results indicate that C-WFPM_0.1_ selectively alters genes related to angiogenesis, cell adhesion, and extracellular matrix (ECM), etc., whereas LS-WFPM_0.1_ selectively impacts ribosomal and translational pathways.

### High molecular weight PAH induces ovarian hyperandrogenism and AhR activation *in vitro*

To validate whether PAHs contained in WFPM_0.1_ underlie the ovarian-disrupting effects observed, *in vitro* cultured follicles were treated with BaP, an established high molecular-weight PAH, and naphthalene (NaP), a low molecular weight PAH, using the same exposure regimen. Follicles from all BaP groups exhibited comparable survival and growth patterns (Figure 11A). Following *in vitro* ovulation on day 6, both vehicle-and BaP-treated follicles ruptured and released MII oocytes with extrusion of the first polar body on day 7 (Figure 11A). Estradiol secretion on day 6 was comparable between groups (Figure 11B). Consistent with the WFPM_0.1_ exposure data, however, follicles exposed to 0.1 µM BaP exhibited a significant increase in testosterone production (Figure 11B). Different from BaP, NaP did not alter any ovarian endpoints (Figure S5).

**Figure 11.**
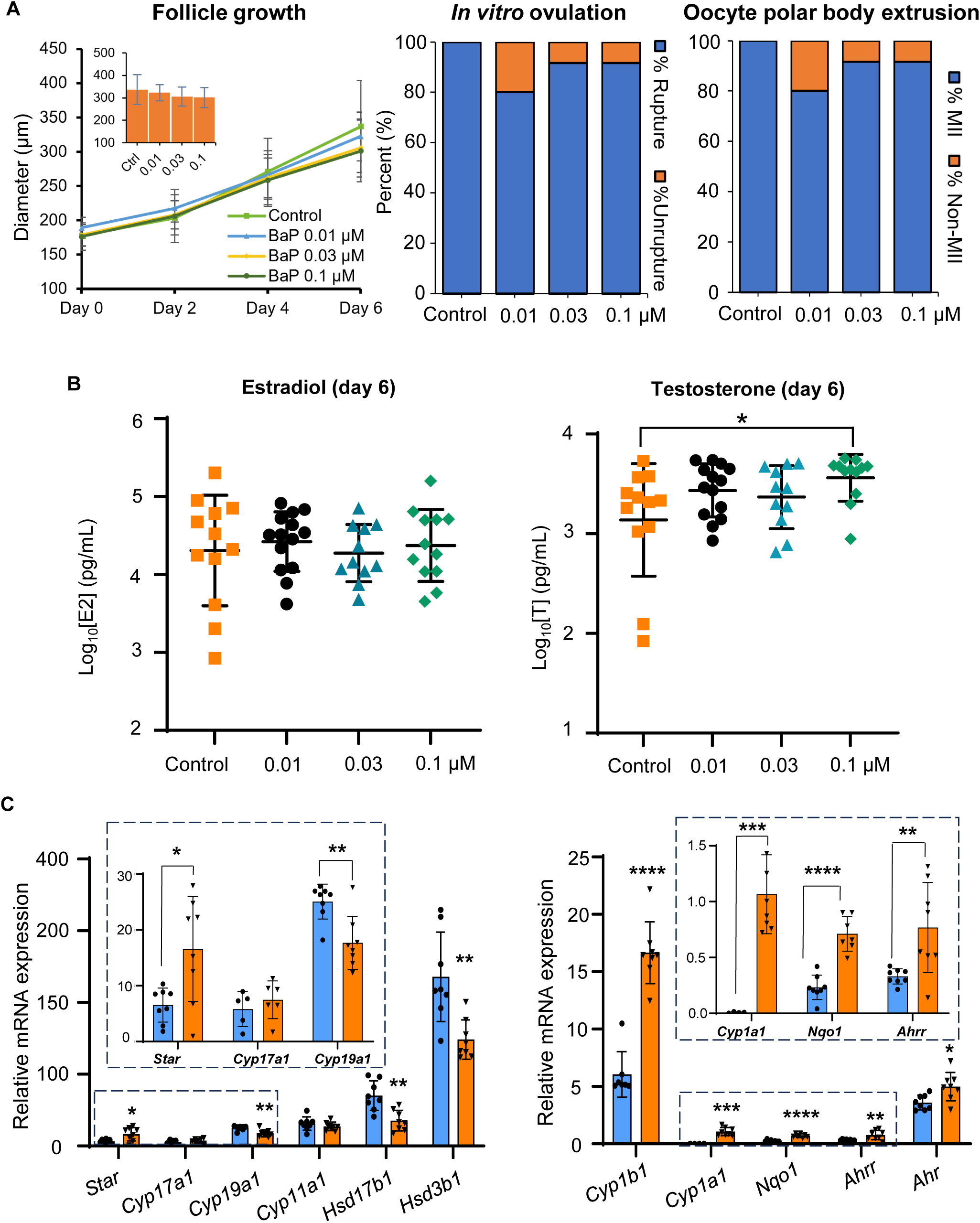
Effects of *in vitro* exposure to BaP during follicle maturation window (day 2-6) on follicle growth, ovulation, hormone secretion and ovarian gene expression. (A) Follicle diameter of control and BaP exposed group (N=10-13 follicles per group) at various concentrations; Percent of ruptured, un-ruptured follicles and ovulated MII oocytes on day 7 of eIVFG. (C) Average Log_10_ concentration (pg/mL) of estradiol and testosterone in the conditioned follicle culture media collected on day 6 of eIVFG (N=11-14 follicles per group). Data were analyzed with one-way ANOVA followed by Dunnett’s multiple comparisons test. Error bars represent mean ± standard deviation. \**p < 0.05.* For *in vitro* ovulation and polar body extrusion in (A), data were analyzed using Fischer’s exact test. (C) Relative mRNA expression of steroidogenic and AhR target genes in control and 0.1 µM BaP treated single follicles (5-8 follicles per group). The mRNA expression levels were normalized by the expression of beta actin (*Actb*). Data was analyzed with Student’s t-test. Shown are mean ± standard deviation; \**p* 0.05, \*\**p < 0.01,* \*\*\**p<0.001,* \*\*\*\**p < 0.0001*.

Follicles treated with vehicle or 0.1 µM BaP were collected on day 6 to examine the expression of steroidogenesis and AhR-related genes by RT-qPCR. Similar to WFPM_0.1_, BaP significantly increased the expression of *Star* and decreased *Cyp19a1*, *Hsd17b1*, and *Hsd3b1* (Figure 11C). Moreover, BaP significantly induced the expressions of AhR targets (*Ahrr, Cyp1a1, Cyp1b1, Nqo1*) and *Ahr* itself (Figure 11C). Taken together, these results suggest that high-molecular-weight PAHs in WFPM_0.1_ are the primary drivers of AhR activation in follicular cells and of ovarian hyperandrogenism.

### AhR inhibition reverses WFPM_0.1_-induced ovarian hyperandrogenism

To further examine the mechanistic causation between WFPM_0.1_-activated AhR and ovarian hyperandrogenism, follicles were co-treated with LS-WFPM_0.1_ and resveratrol, which has been established as an AhR inhibitor [125–127]. The treatment of resveratrol alone did not affect testosterone secretion (Figure 12A). However, resveratrol reversed ovarian hyperandrogenism (Figure 12B) and induction of AhR target genes caused by LS-WFPM_0.1_ (Figure 12C). Taken together, these results indicate that WFPM_0.1_ induces ovarian hyperandrogenism by activating AhR.

**Figure 12.**
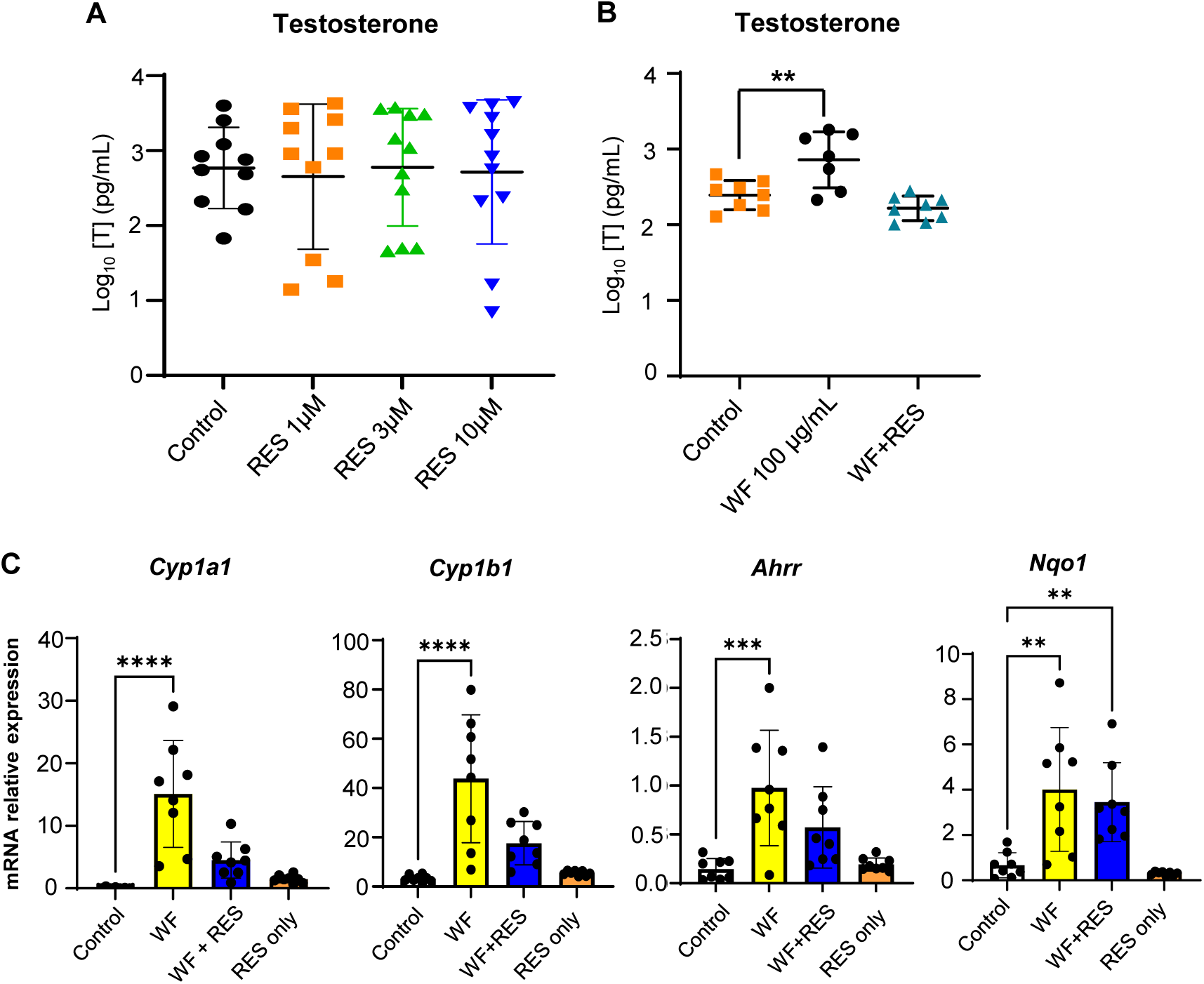
AhR inhibition reverses WFPM_0.1_ induced ovarian hyperandrogenism. (A) Average Log_10_ concentration (pg/mL) of testosterone in the conditioned follicle culture media collected from follicles treated with various concentrations of resveratrol (RES) on day 6 of eIVFG (N=10-11 follicles per group). (B) Average Log_10_ concentration (pg/mL) of testosterone in the conditioned follicle culture media collected from control, or 100 µg/mL LS-WFPM_0.1_ (WF) treated, or 100 µg/mL LS-WFPM_0.1_ and 10 µM resveratrol cotreated (WF+RES) single follicles on day 6 of eIVFG (N=7-8 follicles per group). (C) Relative mRNA expression of major AhR target genes examined by RT-qPCR in control, WF, WF+RES or RES only treated follicles. N=8 follicles per group. Expression levels were normalized by the expression of *Gapdh*. Data were analyzed with one-way ANOVA followed by Dunnett’s multiple comparisons test. Error bars represent mean ± standard deviation. \*\**p < 0.01,* \*\*\**p<0.001,* \*\*\*\**p < 0.0001*.

### BMD modeling for *in vitro* and endpoints

To quantitatively assess the dose-response relationship, we applied Bayesian benchmark dose (BBMD) analysis to testosterone secretion data from follicles exposed to LS-WFPM_0.1_ and C-WFPM_0.1_. The results revealed that the calculated median benchmark concentrations producing a 10% change in testosterone (BMC_10_) are 39.66 for LS-WFPM_0.1_ and 77.36 μg/mL for C-WFPM_0.1_, with BMCL_10_ at 0.396 and 30.89 μg/mL, respectively. These findings indicate that LS-WFPM_0.1_ triggered measurable androgenic effects at lower concentrations than C-WFPM_0.1_.

## Discussion

Wildfires have become more frequent and widespread worldwide [1–6], posing a major concern due to elevated WFPM and its detrimental impact on air quality and health. Here, we employed an *in vivo* intratracheal exposure mouse model and a 3D ovarian follicle culture system, complemented with molecular, transcriptomic, and computational approaches, to demonstrate that WPFM_0.1_ induces ovarian hyperandrogenism through AhR activation.

During severe wildfire events, ambient PM_2.5_ can remain extremely high for several weeks, with mean values exceeding 300-700 µg/m³ over extended periods [11, 128]. The 2023 Canadian wildfires led to ambient PM_2.5_ levels> 400 µg/m³ in New York City, hundreds of miles away from the wildfire origin in Quebec, Canada [13, 14]. Populations living in the fire zone and in areas where WFPM migrates to are experiencing accumulated exposures comparable to our modeled doses. Although intratracheal instillation delivers this burden in a single administration (bolus), the method allows a precise control of the deposited dose and is widely used to mimic real-world pulmonary exposure to PM [129, 130]. The *in vitro* dose ranges were also estimated based on WFPM levels recorded during the 2023 Canadian wildfire [7, 13, 14]. Thus, the WFPM_0.1_ exposure regimen used here represents a high but still environmentally and human-relevant dose, simulating exposure to high levels of WFPM_0.1_ during severe wildfire seasons.

The LS-WFPM_0.1_ used in our study was generated through the complete combustion of pinewood, a major wood in Canadian and California wildfires [8, 9]. Our recent study has demonstrated that the real-world WFPM mostly contained nano-sized particles, reaching a peak of 145.8 μg/m^3^, approximately 10 times higher than the World Health Organization (WHO) Air quality guidelines [7]. These two particle types exhibited both similarities and differences in terms of their physicochemical properties, indicative of their complex chemistry and both contain primarily high levels of organic compounds including high molecular weight PAHs. Total PAH concentration in the size fractionated PM samples was 98.1 ng/m^3^ of which 14.1% was bound to WFPM_0.1_ and 41.3% was bound to WFPM_0.1–2.5_, while the remaining was bound to larger sized particles. WFPM_0.1_ were mainly associated with high molecular weight PAHs which have been shown to bioaccumulate and cause ovarian toxicity [131, 132]. PAH profile of C-WFPM_0.1_ showed similarities as well as differences with LS-WFPM_0.1._ For instance, retene, a well-known marker of wood combustion, was found in both particle types at the highest percentage among all PAHs. They also contained several medium to high-molecular-weight PAHs, including indeno(1,2,3-cd)pyrene, benzo(g,h,i)perylene, and chrysene, which are known AhR agonists [133–135]. However, LS-WFPM_0.1_ uniquely contained benzo[a]pyrene (BaP), a highly carcinogenic PAH and a potent AhR agonist, suggesting subtle compositional differences that might influence the downstream differences in endocrine effects. As expected, the real-world C-WFPM_0.1_ contains other chemical compounds such as heavy metals, which are emitted from other sources, such as traffic-related emissions. Collectively, these findings indicate that although there are distinctions between LS- and C-WFPM_0.1_, primarily due to other sources in real-world PM sampling, LS-WFPM_0.1_ reproduces the key elemental fingerprints of the real-world WFPM_0.1_.

WFPM_0.1_ disrupted ovarian steroidogenesis in both *in vitro* and *in vivo* systems. *In vivo*, intratracheal WFPM_0.1_ exposure slightly elevated serum concentrations of estradiol and testosterone, whereas *in vitro* exposure to both LS and C- WFPM_0.1_ promoted testosterone secretion without affecting estradiol. One possible explanation lies in differential gene expression changes caused by LS-WFPM_0.1_. *In vivo*, we observed a trend toward increased *Star* expression, which promotes cholesterol transport into mitochondria and serves as the first step of steroidogenesis. Increased *Star* expression has been previously linked to increased E2 biosynthesis in granulosa cells [136]. *Star* remained unchanged *in vitro*, consistent with the absence of estradiol elevation. Notably, *Cyp19a1*, a gene encoding aromatase that converts testosterone to estradiol, remained unchanged *in vivo*, which may allow accumulated testosterone to be efficiently converted to estradiol, consistent with the slight increase of estradiol observed *in vivo*. By contrast, *Cyp19a1* was downregulated *in vitro*, limiting the conversion of testosterone to estradiol, which explains why testosterone accumulated *in vitro* without a concomitant increase in estradiol.

Within the *in vitro* model, both LS- and C-WFPM_0.1_ produced broadly similar steroidogenic outcomes, characterized by increased testosterone secretion and suppression of genes involved in estradiol synthesis, including *Cyp19a1* and *Hsd17b1*. However, the two WFPM sources exhibited distinct upstream disruptions within the steroidogenic pathway. WFPM_0.1_ selectively decreased *Cyp17a1*, whereas C-WFPM_0.1_ significantly reduced *Cyp11a1*, the rate-limiting enzyme responsible for initiating steroid biosynthesis. These source-specific differences suggest that while both LS- and C-WFPM_0.1_ converge on androgen accumulation *in vitro*, they may do so through disruption of similar as well as different transcriptional mechanisms, potentially reflecting compositional differences between LS and C-WFPM_0.1_.

The endocrine disrupting effect of WFPM_0.1_ is also manifested by estrous cycle alterations. Mice exposed to WFPM_0.1_ spent significantly less time in proestrus and more time in diestrus, similar to both hyperandrogenic mouse models and mice treated with estrogenic compounds that lead to higher serum E2 levels [137–139]. The slightly elevated estradiol might interfere with the neuroendocrine feedback mechanisms in the pituitary and hypothalamus, creating an endocrine environment in the ovary that suppresses progression from diestrus to proestrus, thereby disrupting estrous cycles. Of course, a central effect of the WFPM_0.1_ cannot be completely ruled out.

Our recent work and those by others demonstrate that both environmental and engineered nanoscale PM, including plastic and metallic nanoparticles (NPs), are able to cross biological barriers in the lung and gut, enter systemic circulation, and reach the placenta and other non-respiratory tissues [140, 141]. Prior studies also reported ovarian accumulation of inhaled PM_0.1_ [42–45]. In mice, inhaled cerium dioxide NPs were present in cumulus cells [42]. In rats, inhaled gold NPs accumulated in granulosa cells [43]. Nano-sized ambient black carbon particles have also been detected in human ovarian tissues and follicular fluid [45]. As a ligand-activated transcription factor, AhR can be activated by diverse xenobiotics, including dioxin and PAHs [142–144]. Upon ligand binding, AhR heterodimerizes with AhR nuclear translocator (ARNT) and binds to xenobiotic response elements (XREs) to induce target genes, such as *Cyp1a1*, *Cy1b1*, *Ahrr*, and *Nrf2* [119, 145, 146]. Although we did not directly quantify WFPM_0.1_ in the ovary, the consistent induction of AhR target genes in follicles, together with hyperandrogenism observed both *in vivo* and *in vitro*, strongly suggest the ovarian accumulation and direct effects of WFPM_0.1_. In future studies, toxicokinetic experiments will be used to quantify the ovarian accumulation of PAHs and WFPM.

The induction of AhR target genes and rescue of WFPM_0.1_ or BaP-induced testosterone overproduction by AhR antagonist suggest that PAHs in WFPM_0.1_ activate AhR, thereby resulting in hyperandrogenism. Prior studies have shown that the induction of phase I enzymes of CYP1A1 and CYP1B1 can metabolize parent compounds into reactive intermediates and generate electrophiles and reactive oxygen species (ROS), leading to oxidative stress [147–151]. Consistent with this, both LS- and C-WFPM_0.1_, as well as BaP, upregulated CYP1 genes and genes related to oxidative stress. In addition to AhR-dependent mechanisms, ultrafine particles themselves are well-established inducers of oxidative stress due to their high surface area to volume ratio, redox-active organic compounds and associated transition metals, which can directly generate ROS through surface chemistry and cellular interactions, leading to mitochondrial damage [152–154].

Elevated ROS have been detected in the serum and follicular fluid of PCOS women with hyperandrogenism [155], and ROS modulate MAPK signaling and enhance CYP17A1 activity, a key enzyme in androgen biosynthesis [156]. Beyond oxidative stress, AhR also interacts with SF-1 (also called NR5A1) [157], a transcription factor that regulates *Star* expression [158]. Here, we propose that WFPM_0.1_ may promote androgen synthesis by inducing oxidative stress, either AhR-dependent or-independent, and/or by modulating AhR-mediated expression of steroidogenic genes. Further mechanistic studies are required to deeply dive into these pathways and define the specific role of AhR in mediating WFPM_0.1_–induced ovarian hyperandrogenism.

Resveratrol has been characterized as a natural AhR antagonist that competitively binds to AhR [125, 127]. We demonstrated that resveratrol reversed WFPM_0.1_-induced hyperandrogenism and effects on AhR-target genes. Resveratrol is also an antioxidant [159], which might be mediated through the activation of sirtuin 1 (SIRT1), a redox-sensitive deacetylase that improves mitochondrial function [160], and/or through activating Nrf2 to induce antioxidant enzymes [161]. Clinically, resveratrol has been shown to reduce androgen secretion and improve menstrual cyclicity in PCOS patients with hyperandrogenism and elevated ROS [162, 163]. These findings suggest that resveratrol may attenuate WFPM_0.1_-induced testosterone overproduction as an AhR antagonist and/or as an antioxidant.

Nrf2 is a central transcription factor against xenobiotic exposure and oxidative stress [147, 164]. Activated AhR upregulates Nrf2 which heterodimerizes with small musculoaponeurotic fibrosarcoma (sMAF) protein and binds to the antioxidant response element (ARE), inducing antioxidant enzymes including SOD, GSR, and phase II detoxication enzymes such as NQO1 and GST [164, 165]. In our study, ovarian follicles exposed to either LS- or C-WFPM_0.1_ showed elevated expression of *Nrf2* and downstream antioxidant genes, including *Sod1*, *Sod2*, *Gstp1*, *Gsr*, and *Nqo1*. However, WFPM_0.1_-treated follicles still exhibited hyperandrogenism, suggesting that although NRF2 activation provides a compensatory defense, it is insufficient to counteract excessive ROS generated from the activated AhR signaling. Moreover, excessive ROS may perturb redox-sensitive transcriptional networks in follicular cells. For instance, *Nqo1* and *Gstp1* regulate intracellular NADPH and GSH pools, which serve as critical cofactors for steroidogenic enzymes. Thus, increased *Nrf2* may shift redox flux in ways that inadvertently modulate steroidogenesis. In line with this, our data showed upregulation of *Nqo1*, *Gstp1*, *Sod1/2*, *Gsr* following WFPM_0.1_ exposure, suggesting that the Nrf2-driven antioxidant responses may shift NADPH and GSH availability away from balanced steroidogenesis, thereby promoting androgen accumulation.

Both WFPM types altered a common set of AhR target genes and increased testosterone in exposed follicles, indicating that the overlapping toxic components, particularly PAHs, result in similar ovarian toxicities observed. At the transcriptomic level, LS- and C-WFPM_0.1_ showed overlapping as well as distinct pathway perturbations. Approximately 25% of the DEGs were shared between the treatments of LS- and C-WFPM_0.1_, representing common cellular responses to WFPM_0.1_ exposure. These common transcriptomic changes converge on processes that regulate cytoskeletal organization, transcriptional control, protein turnover, and stress signaling. Cytoskeletal remodeling can influence intracellular trafficking and organelle positioning, including mitochondria and endoplasmic reticulum (ER), which are essential sites for steroid hormone biosynthesis. Concurrently, altered transcriptional coregulator activity and protein turnover pathways can modify the abundance, stability, and activity of key steroidogenic enzymes. Together, these coordinated perturbations may increase testosterone production directly, or create a cellular milieu favoring androgen accumulation. In addition, there were distinct DEGs and signaling pathways between the two particle types. LS-WFPM_0.1_ uniquely altered 3,453 genes that are associated with a primary disruption of translational and protein-quality-control machinery that might cause steroidogenic dysregulation without markedly affecting luteal progesterone production. By contrast, C-WFPM_0.1_ selectively changed 2,361 genes enriched for cytoskeletal remodeling, cell–ECM adhesion and angiogenesis-related signaling pathways. These pathways are critical for CL formation and function. Disruption in angiogenic signaling can impair CL development and function, leading to reduced progesterone secretion [166, 167]. Additionally, extensive ECM remodeling is required for successful luteinization [168]. The selective alteration of these pathways by C-WFPM_0.1_, but not LS-WFPM_0.1_, aligns with our observation that only C-WFPM_0.1_ decreased progesterone secretion in formed CL. This suggests that C-WFPM_0.1_ may contain additional bioactive constituents beyond PAHs that lead to distinct ovarian toxicities.

Our RNA-seq data revealed distinct but converging transcriptomic changes in response to the treatment of LS-WFPM_0.1_ *in vivo* and *in vitro* and C-WFPM_0.1_ *in vitro*. *In vivo* data highlighted enrichment of “Cholesterol metabolism” and “IL-17 signaling”, indicating disruptions in ovarian steroidogenesis and inflammatory pathways. Additional enrichment of “Tight junctions”, “Cell-substrate adhesion”, and ECM-related GO terms suggested impaired follicular architecture and follicular cell interactions. Gap junctions between granulosa cells and oocytes depend on ECM integrity to sustain direct intercellular communication, nutrient exchange, normal folliculogenesis, and steroidogenesis [169, 170]. Disruption of these networks may underlie high testosterone secretion following WFPM_0.1_ exposure. In follicles treated with LS-WFPM_0.1_, DEGs were enriched in pathways related to neurodegeneration, “Mitophagy”, “Autophagy”, and “Proteosome”. Although these annotations reflect the neurological contexts, a shared mechanism is mitochondrial dysfunction and elevated ROS [171, 172]. Given that granulosa cells rely on mitochondria and ATP supply to support follicle/oocyte health, mitophagy enrichment may suggest mitochondrial damage [173]. Follicles treated with C-WFPM_0.1_ had DEGs related to inflammation and stress, including “AGE-RAGE signaling pathway”, “cellular senescence”, “focal adhesion”, “Rap1 signaling”, and multiple GO terms involved in ECM/actin filament organization. As a hallmark of oxidative stress, AGE-RAGE can activate NF-κB-mediated inflammation, a key feature implicated in PCOS [174]. Dysregulated focal adhesion and ECM organization are also shown to disrupt follicular development and hormone secretion [169], including PCOS ovaries with excess collagen deposition and fibrosis [175–177]. These findings suggest that WFPM_0.1_ disrupts both biochemical (hormonal/redox) and biomechanical (ECM/adhesion) environments of follicles, converging to promote androgen excess.

There is currently limited epidemiological data regarding the effect of WFPM on oocyte health. A recent retrospective cohort study correlated decreased gamete quality and blastocyst yield with a severe wildfire event in Oregon [178]. However, there was a limited sample size to show statistical power and there was no WFPM data at the time of data collection. Our results indicate that oocytes from WFPM-treated follicles were morphologically similar to controls. However, single-oocyte RNA-seq analysis discovered 5 significantly altered MEGs. Among 3 up-regulated MEGs, *Khdc3* and *Zbed3* encode key components of the subcortical maternal complex (SCMC), a multiprotein complex selectively expressed in oocytes and early embryos, which plays crucial roles in early embryo development [179]. The third up-regulated gene, *Mapk3*, encodes mitogen-activated protein kinase 3 (MAPK3, also known as extracellular signal-regulated kinase 1 or ERK1), which is essential for oocyte maturation [180]. Of the two down-regulated genes, *Dppa3* (also known as *Stella* or *Pgc7*) has been established to express in primordial germ cells (PGCs), oocytes, and preimplantation embryos [181]. It facilitates DNA demethylation by directing binding to RING finger–type E3 ubiquitin ligase UHRF1 [182]. While *Dppa3^-^*^/-^ female mice have normal folliculogenesis, embryos fail to progress to the blastocyst stage [183]. The other down-regulated gene, *Ago2*, encodes Argonaute 2 (AGO2), a key component of the RNA-induced silencing complex (RISC). AGO2 mediates gene repression by targeting microRNAs (miRNAs) or small interfering RNAs (siRNAs) [184]. It has been reported that *Ago2* is expressed in oocytes and pre-implantation embryos [185], and mice with global deletion of *Ago2* exhibit early embryonic lethality [186]. Moreover, although female mice with selective deletion of *Ago2* in oocytes have normal follicle development and ovulation, they are infertile and ovulate oocytes with defective spindle and chromosome alignment [185, 187]. These findings highlight the crucial roles of these MEGs in supporting oocyte developmental competence, and exposure to WFPM may alter MEGs and impair oocyte quality, leading to defective reproductive outcomes

While PAHs are potent AhR agonists, the additional constituents of WFPM_0.1_ may also contribute to ovarian toxicity through AhR-dependent and/or -independent mechanisms. For instance, brominated flame retardants have been shown to activate AhR [188] and disrupt ovarian steroidogenesis in rat ovaries [189]. Phthalates and their metabolites can disrupt ovarian steroidogenesis via non-AhR pathways [190]. Therefore, WFPM_0.1_-induced ovarian hyperandrogenism might not be solely attributable to PAHs but could also involve other WFPM_0.1_ constituents, either by acting as an AhR agonist or through non-AhR pathways.

Our WFPM_0.1_ results are concordant with reports on other types of PM such as the traffic-related ultrafine particles including the diesel exhaust particles (DEPs) that contain a majority of ultrafine particles by number [191]. DEPs are well documented to also contain high levels of PAHs and activate AhR signaling and to generate reactive metabolites and ROS in exposed tissues and cells, making CYP1A1 a sensitive biomarker of combustion-derived particle exposure [192, 193]. Many other ambient PM_2.5_ inhalation studies have also been linked to ovarian dysfunction in rodents, including increased follicular atresia, granulosa cell apoptosis, altered steroidogenic gene expression, and inflammation [194–196].

In summary, our study reveals that exposure to WFPM_0.1_ interferes with ovarian steroidogenesis, induces ovarian hyperandrogenism, and alters the oocyte transcriptome and key MEGs, possibly through the activation of AhR in follicular somatic cells by PAHs contained in WFPM_0.1_. Hyperandrogenism is a central feature of PCOS and has a wide range of consequences for women’s overall health. Clinically, androgen excess manifests as hirsutism, acne, and ovulatory dysfunction, and is a major contributor to infertility in affected women [197]. Hyperandrogenism is also tightly linked to pregnancy complications, cardiovascular diseases, and metabolic syndrome including insulin resistance, dyslipidemia, and impaired glucose tolerance [197, 198]. These reproductive and metabolic disturbances also coincide with a higher prevalence of anxiety, depression, and diminished quality of life among women with PCOS [199]. Given more frequent and intense wildfires in the US and worldwide and the ubiquitous exposure of populations to WFPM, our results can be used to tailor guidance to protect women’s reproductive health and fertility. Our results highlight that WFPM is an important and currently understudied exposure that may detrimentally impact female reproductive function. Given more frequent and intense wildfires in the US and worldwide and the ubiquitous exposure of populations to WFPM, additional research on this topic is warranted to protect women’s reproductive health and fertility from this emerging threat.

## Authors’ roles

K. Mali and D. Zhang contributed equally to the experimental design, data collection, analysis, interpretation, and manuscript writing. L. Bazina and K. Moularas contributed to LS-WFPM synthesis, preparation, physicochemical characterization, and related data collection, result interpretation, and manuscript writing. E. Abramova and A. Gow contributed to the intratracheal instillation-related experimental design and conduction. J. Zhang contributed to bioinformatic data analysis and related manuscript writing. T. Zhan and P. Pattarawat partially contributed to *in vivo* and *in vitro* exposure experiments and data collection. Q. Zhang contributed to BMD modeling and manuscript editing. A.J. Gaskins contributed to manuscript editing. P. Demokritou and S. Xiao conceived of the project, designed experiments, analyzed and interpreted data, wrote the manuscript, and provided final approval of the manuscript.

## Funding and acknowledgments

This work was supported by the National Institutes of Health (NIH) R01ES032144, R01ES037796 and P30ES005022 to S. Xiao; 1R01ES037796-01, 1R21ES038066-01 and National Science Foundation (NSF) 2524102 to P. Demokritou.

