## Supplementary material for "Wildfire emitted particulate matter induces ovarian hyperandrogenism through aryl hydrocarbon receptor activation": Fig S1

### Slide 1
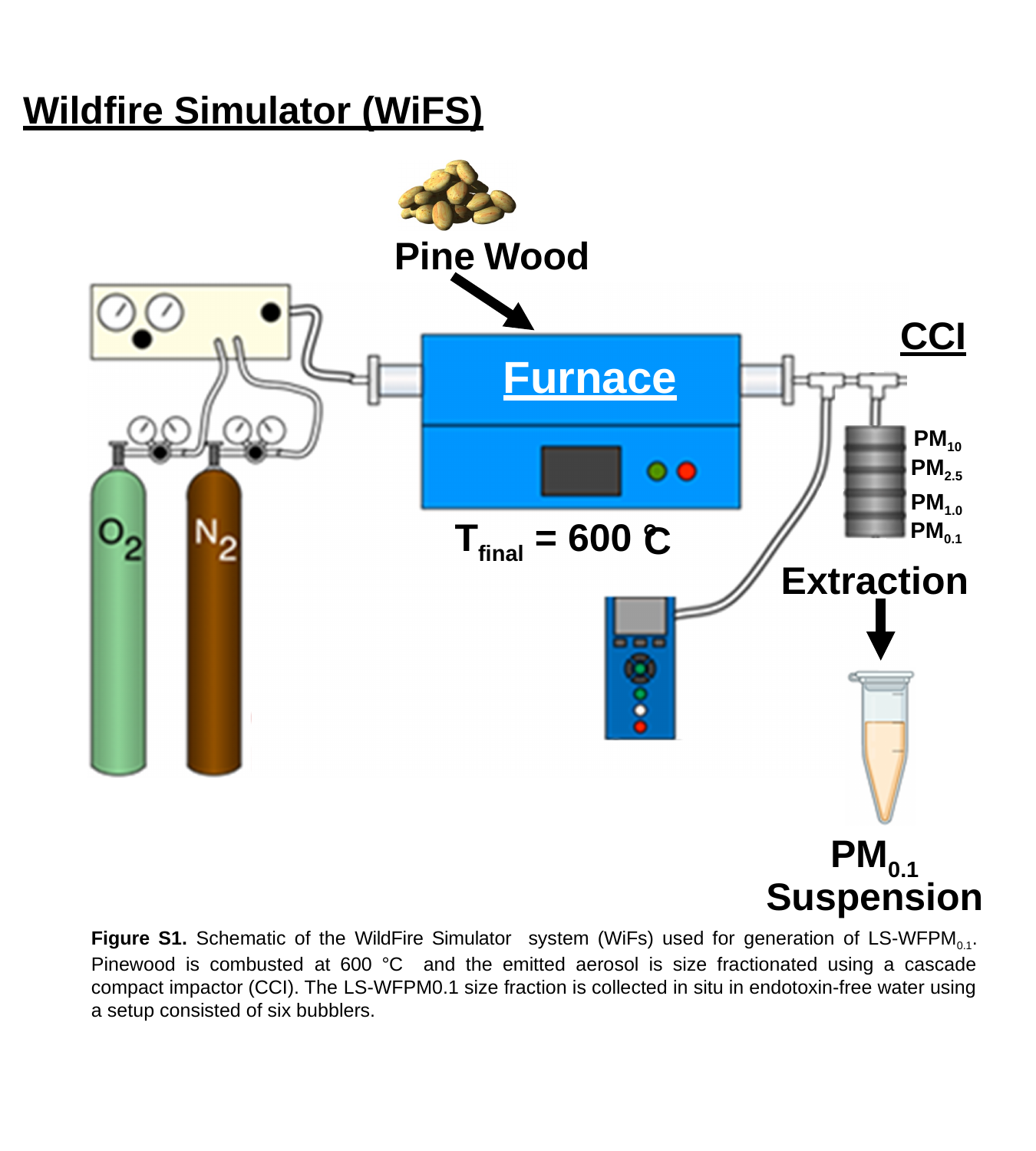

Wildfire Simulator (WiFS)
Pine Wood
CCI
Furnace
PM_10_
PM_2.5_
PM_1.0_
T_final_ = 600 °
C
PM_0.1_
Extraction
PM_0.1_
Suspension
Figure S1. Schematic of the WildFire Simulator system (WiFs) used for generation of LS-WFPM_0.1_. Pinewood is combusted at 600 °C and the emitted aerosol is size fractionated using a cascade compact impactor (CCI). The LS-WFPM0.1 size fraction is collected in situ in endotoxin-free water using a setup consisted of six bubblers.
PM_10_
PM_2.5_
PM_1.0_
PM_0.1_
T_final_ = 600 °
C
Extraction
PM_0.1_
Suspension
