## Supplementary material for "Wildfire emitted particulate matter induces ovarian hyperandrogenism through aryl hydrocarbon receptor activation": Fig S2

### Slide 1
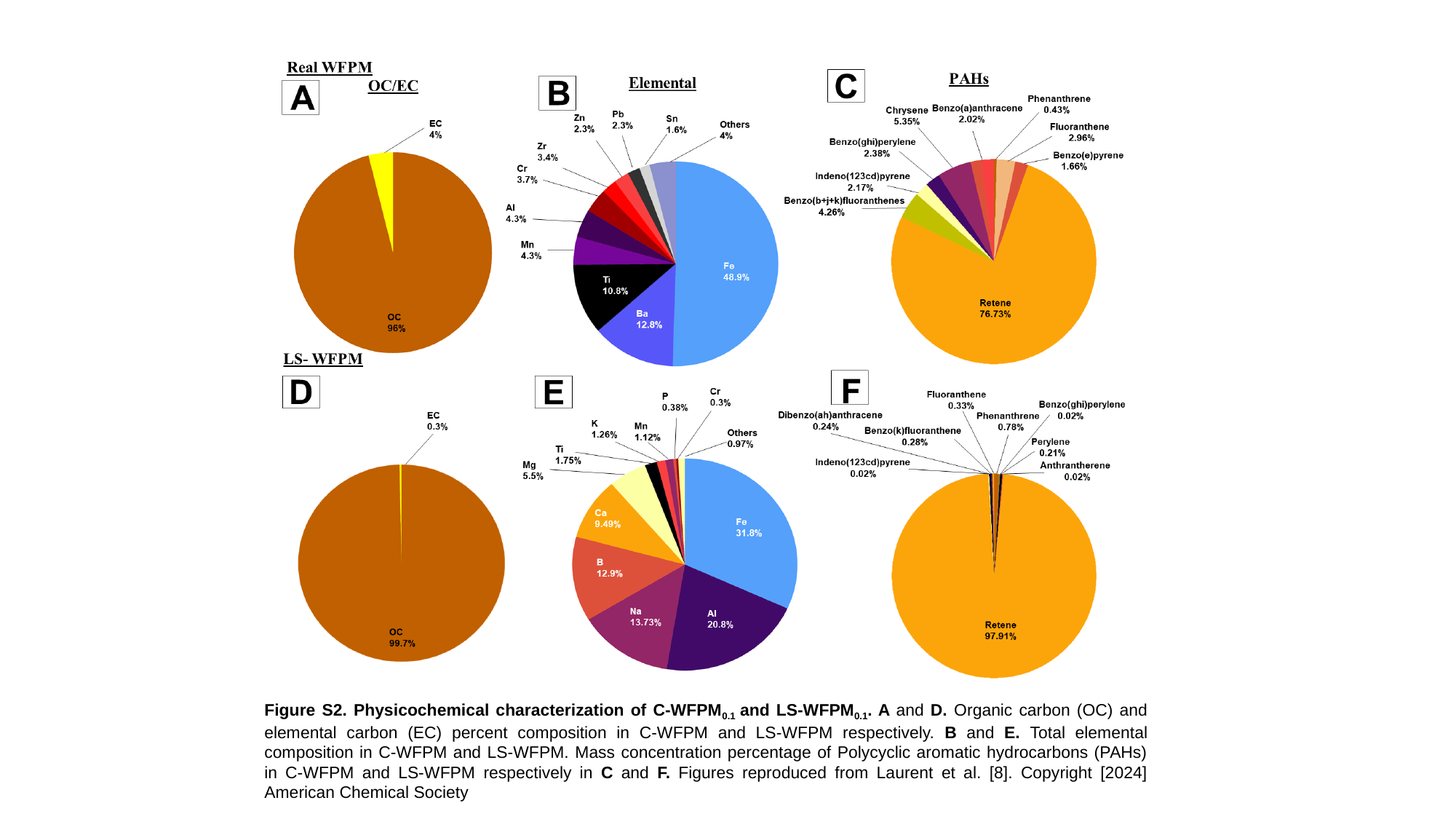

Figure S2. Physicochemical characterization of C-WFPM_0.1_ and LS-WFPM_0.1_. A and D. Organic carbon (OC) and elemental carbon (EC) percent composition in C-WFPM and LS-WFPM respectively. B and E. Total elemental composition in C-WFPM and LS-WFPM. Mass concentration percentage of Polycyclic aromatic hydrocarbons (PAHs) in C-WFPM and LS-WFPM respectively in C and F. Figures reproduced from Laurent et al. [8]. Copyright [2024] American Chemical Society
