## Supplementary figures and images for "Wildfire emitted particulate matter induces ovarian hyperandrogenism through aryl hydrocarbon receptor activation"

### Fig S3

#### Slide 1
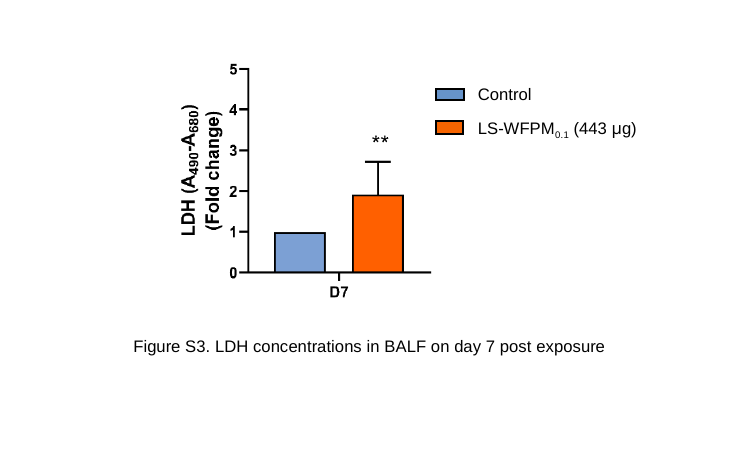

Control
LS-WFPM_0.1_ (443 µg)
Figure S3. LDH concentrations in BALF on day 7 post exposure
