## Supplementary material for "Wildfire emitted particulate matter induces ovarian hyperandrogenism through aryl hydrocarbon receptor activation": Fig S4

### Slide 1
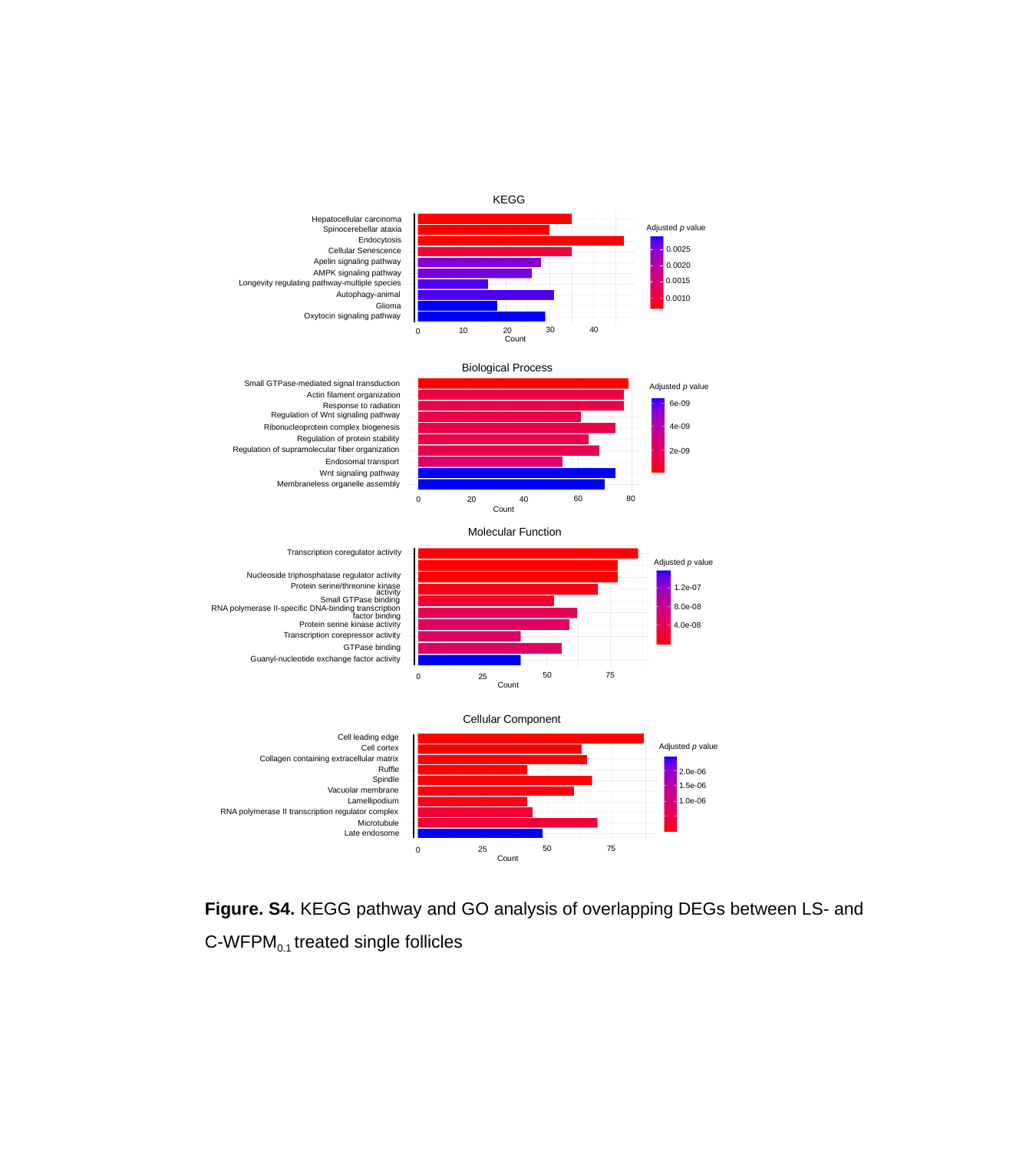

KEGG
Hepatocellular carcinoma
Spinocerebellar ataxia
Cellular Senescence
Apelin signaling pathway
AMPK signaling pathway
Longevity regulating pathway-multiple species
Autophagy-animal
Oxytocin signaling pathway
Endocytosis
Glioma
30
40
10
20
0
Count
Adjusted p value
0.0025
0.0020
0.0015
0.0010
Biological Process
Small GTPase-mediated signal transduction
Actin filament organization
Response to radiation
Regulation of Wnt signaling pathway
Regulation of protein stability
Endosomal transport
Wnt signaling pathway
Membraneless organelle assembly
Ribonucleoprotein complex biogenesis
Regulation of supramolecular fiber organization
80
60
0
20
40
Count
Adjusted p value
6e-09
4e-09
2e-09
Molecular Function
Transcription coregulator activity
Nucleoside triphosphatase regulator activity
Protein serine/threonine kinase activity
Small GTPase binding
RNA polymerase II-specific DNA-binding transcription factor binding
Protein serine kinase activity
Transcription corepressor activity
GTPase binding
Guanyl-nucleotide exchange factor activity
Adjusted p value
1.2e-07
8.0e-08
4.0e-08
75
50
25
0
Count
Cellular Component
Cell leading edge
Cell cortex
Collagen containing extracellular matrix
Ruffle
Spindle
Vacuolar membrane
Lamellipodium
RNA polymerase II transcription regulator complex
Microtubule
Late endosome
75
50
25
0
Count
Adjusted p value
2.0e-06
1.5e-06
1.0e-06
Figure. S4. KEGG pathway and GO analysis of overlapping DEGs between LS- and C-WFPM_0.1_ treated single follicles
