## Supplementary material for "Wildfire emitted particulate matter induces ovarian hyperandrogenism through aryl hydrocarbon receptor activation": Fig S5

### Slide 1
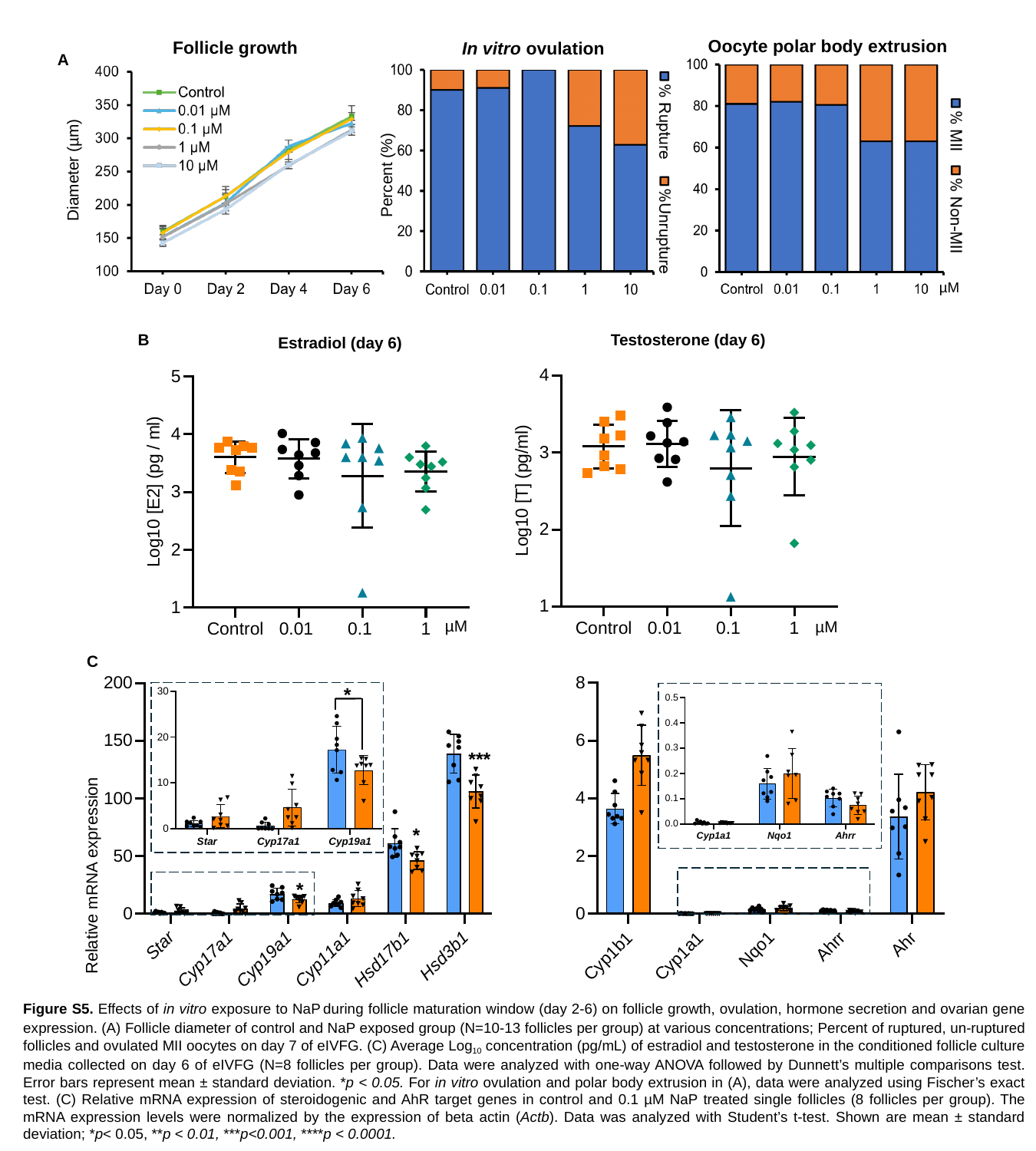

Oocyte polar body extrusion
Follicle growth
In vitro ovulation
A
Diameter (µm)
Percent (%)
 µM
Testosterone (day 6)
B
Estradiol (day 6)
µM
µM
C
*
***
*
Relative mRNA expression
*
Figure S5. Effects of in vitro exposure to NaP during follicle maturation window (day 2-6) on follicle growth, ovulation, hormone secretion and ovarian gene expression. (A) Follicle diameter of control and NaP exposed group (N=10-13 follicles per group) at various concentrations; Percent of ruptured, un-ruptured follicles and ovulated MII oocytes on day 7 of eIVFG. (C) Average Log_10_ concentration (pg/mL) of estradiol and testosterone in the conditioned follicle culture media collected on day 6 of eIVFG (N=8 follicles per group). Data were analyzed with one-way ANOVA followed by Dunnett’s multiple comparisons test. Error bars represent mean ± standard deviation. *p < 0.05. For in vitro ovulation and polar body extrusion in (A), data were analyzed using Fischer’s exact test. (C) Relative mRNA expression of steroidogenic and AhR target genes in control and 0.1 µM NaP treated single follicles (8 follicles per group). The mRNA expression levels were normalized by the expression of beta actin (Actb). Data was analyzed with Student’s t-test. Shown are mean ± standard deviation; *p˂ 0.05, **p < 0.01, ***p<0.001, ****p < 0.0001.
