## Supplementary material for "Wildfire emitted particulate matter induces ovarian hyperandrogenism through aryl hydrocarbon receptor activation": Table S1

| **Gene Symbol** | **Full Name** | **Major functions** | **References** |
| --- | --- | --- | --- |
| *Cyp1a1* | Cytochrome P450, family 1, subfamily a, polypeptide 1 | Encodes CYP1A1, a phase I xenobiotic metabolizing enzyme, catalyzes the oxidative metabolism of AhR ligands like polycyclic aromatic hydrocarbons (PAHs) and halogenated aromatic hydrocarbons (HAHs) | [1, 2] |
| *Cyp1b1* | Cytochrome P450, family 1, subfamily b, polypeptide 1 | Encodes CYP1B1, also metabolizes xenobiotics like PAHs and HAHs; metabolizes 17β-estradiol; involved in retinoic acid metabolism in humans | [1, 3, 4] |
| AhR | Aryl hydrocarbon receptor | Ligand-activated transcription factor regulating xenobiotic metabolism; mediates responses to environmental pollutants (e.g., PAHs, HAHs, dioxins) by activating the induction of genes related to phase I metabolism; plays a role in organ development and function, immune cell differentiation, regulates normal female reproductive functions and fertility | [5, 6] |
| *Ahrr* | Aryl hydrocarbon receptor repressor | Transcriptional repressor that competes with AhR for ARNT binding, providing negative feedback regulation of AhR signaling. | [7, 8] |
| *Nrf2* (also known as *Nfe2l2*) | Nuclear factor, erythroid derived 2, like 2 | Master regulator of antioxidant response; induces phase II metabolizing enzymes like glutathione S-transferase (GST), NAD(P)H:quinone oxidoreductase 1 (NQO1) under oxidative stress. | [2, 9, 10] |
| *Nqo1* | NAD(P)H quinone oxidoreductase 1 | Phase II detoxifying enzyme that reduces quinones and other organic compounds preventing oxidative stress; major target of *Nrf2* gene. | [11, 12] |

**Table S1.** All major AhR related genes and functions.
