## Supplementary material for "Wildfire emitted particulate matter induces ovarian hyperandrogenism through aryl hydrocarbon receptor activation": Table S8

| **Gene Symbol** | **Full Name** | **Fold Change** | **Major functions** | **References** |
| --- | --- | --- | --- | --- |
| *Zbed3* | Zinc finger, BED type containing 3 | 2.54 | In somatic cells, *Zbed3* is known to activate the Wnt/β-catenin signaling pathway. In the oocyte and early embryo, it forms an integral component of the subcortical maternal complex (SCMC) and functions to regulate cytoskeletal dynamics during early embryo development. | [1, 2] |
| *Khdc3,* also known as *Filia* | KH domain containing 3, subcortical maternal complex member | 2.40 | Also forms a critical part of SCMC and functions to maintain spindle assembly, cell cycle check points during early embryogenesis | [3] |
| *Mapk3*, also known as *Erk1* | Mitogen-activated protein kinase 3 | 2.50 | Forms major component of the MAPK/ERK signaling pathway, regulating major cellular processes like proliferation, differentiation and apoptosis; required for oocyte meiotic maturation and pronucleus formation | [4, 5] |
| *Dappa3*, also known as *Stella,* *Pgc7* | Developmental pluripotency-associated 3 | 0.35 | Involved in early embryo viability; protects maternal DNA from demethylation during epigenetic reprogramming in preimplantation zygote | [6] |
| *Ago2* | Argonaute RISC catalytic subunit 2 | 0.45 | Forms core component of RNA-induced silencing complex (RISC) that regulates gene expression through small RNA; maintains oocyte competence through same mechanism | [7] |

**Table S8.** Maternal effect genes (MEGs) in the DEG list from single oocyte SMART-Seq2 RNA seq
