## Supplementary material for "Wildfire emitted particulate matter induces ovarian hyperandrogenism through aryl hydrocarbon receptor activation": Table S11

**Table S11**. Aerosol mass concentration, Mass Median Aerodynamic Diameter, MMAD, and effective density, *ρ*, used as input parameters for lung deposition calculations by MPPD software.

| Particle size fraction | Mass concentration^*^, μg/m^3^ | Mass Median Aerodynamic Diameter, MMAD, µm** | Density^***^, g/cm^3^ |
| --- | --- | --- | --- |
| PM0.1 | 165 | 0.05 | 0.955 |

^*^ Mass concentration was obtained from aerosol mass-size distribution and concentration of C-WFPM using multistage Compact Cascade Impactors (CCIs) reported by Laurent et al. [1].

** Additional MPPD parameters used were specified by Lizonova et al.[2] within the framework of the Yeh–Schum symmetric model [3]. In this configuration, the functional residual capacity was set at 3,300, while the head compartment was represented with a volume of 50 mL. Breathing dynamics assumed nasal respiration at 12 breaths per minute, a tidal volume of 625 mL, and an inspiratory fraction fixed at 0.5 [4]. For both WFPM fractions, the Geometric Standard Deviation (GSD) was assigned a value of 1, indicating a completely uniform distribution of particle sizes.

^***^ Effective density was calculated according to the value previously reported by Leskin et al.[5].
