## Supplementary material for "Wildfire emitted particulate matter induces ovarian hyperandrogenism through aryl hydrocarbon receptor activation": Table S12

**Table S12.** Colloidal characterization of C-WFPM and LS-WFPM in water and cell culture medium at DSEcr (hydrodynamic diameter (dH), polydispersity index (PdI), zeta potential (ζ), specific conductance (σ), and mean effective density (𝝆EV)).

|  | Time (h) | Dispersion | Intensity weighted d_H_ (nm) | PdI | ζ (mV) | σ (mS/cm) |
| --- | --- | --- | --- | --- | --- | --- |
| C-WFPM | 0 | DI WATER | 284.7 ± 71.4 | 0.368 ± 0.007 | -34.3 ± 0.3* | 0.23 ± 0.004* |
|  | 0 | Complete media | 314.5 ± 9.0 | 0.272 ± 0.010 | -8.14 ± 0.44 | 14.8 ± 0.86 |
|  | 24 | Complete media | 351.0 ± 204.4 | 0.961 ± 0.04 |  |  |
| LS-WFPM | 0 | DI WATER | 240.0 ± 4.6 | 0.103 ± 0.023 | -27.40 ± 11.80* | 0.072 ± 0.001* |
|  | 0 | Complete media | 292.2 ± 1.3 | 0.266 ± 0.004 | -9.86 ± 0.70 | 13.2 ± 1.01 |
|  | 24 | Complete media | 293.0 ± 14.2 | 0.270 ± 0.010 |  |  |

* The zeta potential (ζ) and conductivity (σ) values were obtained from previously reported measurements by Bazina et al. (2025) for C-WFPM and by Deloid et al. for LS-WFPM.
