## Supplementary material for "Wildfire emitted particulate matter induces ovarian hyperandrogenism through aryl hydrocarbon receptor activation": Table S13

**Table S13**. Mice mRNA primer sequences used for RT-qPCR

|  | **Gene Name** | **F/R** | **Primer Sequence (5’ to 3’)** |
| --- | --- | --- | --- |
| 1. | *Gapdh* | F | CATCACTGCCACCCAGAAGACTG |
|  |  | R | ATGCCAGTGAGCTTCCCGTTCAG |
| 2. | *Actb* | F | GTGACGTTGACATCCGTAAAGA |
|  |  | R | GCCGGACTCATCGTACTCC |
| 3. | *Star* | F | TTGGGCATACTCAACAACCA |
|  |  | R | CCTTGACATTTGGGTTCCAC |
| 4. | *Cyp11a1* | F | TCAAAGCCAGCATCAAGGAGA |
|  |  | R | TGGCAAAGCTAGCCACCTGTA |
| 5. | *Hsd3b1* | F | AGTGATGGAAAAAGGGCAGGT |
|  |  | R | GCAAGTTTGTGAGTGGGTTAG |
| 6. | *Cyp17a1* | F | CCAGGACCCAAGTGTGTTCT |
|  |  | R | CCTGATACGAAGCACTTCTCG |
| 7. | *Cyp19a1* | F | CATGGTCCCGGAAACTGTGA |
|  |  | R | GTAGTAGTTGCAGGCACTTC |
| 8. | *Hsd17b1* | F | ACTGTGCCAGCAAGTTTGCG |
|  |  | R | AAGCGGTTCGTGGAGAAGTAG |
| 9. | *Ahr* | F | ACTTCACACCTATTGGTTGT |
|  |  | R | ATGCCACTTTCTCCAGTCTT |
| 10. | *Ahrr* | F | GTTGGATCCTGTAGGGAGCA |
|  |  | R | AGTCCAGAGGCTCACGCTTA |
| 11. | *Cyp1a1* | F | TCTCGTGGAGCCTCATGTACCT |
|  |  | R | TGCCGATCTCTGCCAATCA |
| 12. | *Cyp1b1* | F | TTGACCCCATAGGAAACTGC |
|  |  | R | GCTGTCTCTTGGTAGGAGGA |
| 13. | *Cyp1a2* | F | CGTTAGGCCATGTCACAAGTAGC |
|  |  | R | CATCACAAGTGCCCTGTTCAAGC |
| 14. | *Nqo1* | F | AGGCTGCTTGGAGCAAAATA |
|  |  | R | GGCAAATCCTGCTACGAGCACT |
| 15. | *Nrf2* | F | CAGCATAGAGCAGGACATGGAG |
|  |  | R | GAACAGCGGTAGTATCAGCCAG |
| 16. | *Sod1* | F | ATGAGGTCCTGCACTGGTACAG |
|  |  | R | ATCCAGAGAGGAATGAGTGGCG |
