## Supplementary Information for "Wildfire emitted particulate matter induces ovarian hyperandrogenism through aryl hydrocarbon receptor activation"

**Methods**

**Real time monitoring and physicochemical analysis of emitted WFPM_0.1_**

Briefly, emissions from WFPM_0.1_, were characterized using two particle sizing instruments as reported by Laurent et al. [1]. A Scanning Mobility Particle Sizer (SMPS, Model 3080, TSI Inc., Shoreview, MN) was employed to quantify particle number concentrations in the nanoscale range (5–300 nm), while an Aerodynamic Particle Sizer (APS, Model 3321, TSI Inc., Shoreview, MN) was used to measure concentrations of larger particles between 0.5 and 20 µm [1]. To ensure that aerosol concentrations were within the detection limits of the SMPS, the emissions were diluted 200-fold using a Rotating Disk Thermodiluter (Model 379020A, TSI Inc., Shoreview, MN), following the procedure outlined by Sotiriou et al. (2015) [2]. Data acquisition was performed continuously during combustion, with the SMPS recording every 2 minutes and the APS every 10 seconds. Reported SMPS concentrations were subsequently corrected for the dilution factor.

C-WFPM and LS-WFPM were fractionated by size using Harvard Compact Cascade Impactors (HCCIs), separating particles into PM_0.1_ (aerodynamic diameter (dae) < 0.1 µm), PM0.1–2.5 (0.1 ≤ dae < 2.5 µm), PM2.5–10 (2.5 ≤ dae ≤ 10 µm), and PM10 (dae > 10 µm). These fractions were used to determine particle mass size distributions and concentrations, following methodologies described previously [2, 3]. Speciated polycyclic aromatic hydrocarbons (PAHs) and oxygenated PAHs were measured by gas chromatography–mass spectrometry (GC/MS), as outlined in earlier investigations by the research group [1, 4].

**Analysis of elemental and organic carbon and PAH**

Elemental and organic carbon (EC–OC) assessment of the WFPM_0.1_ samples was completed as reported in our recent publication [1]. In brief, one cm^2^ punchout areas of the filters were created and analyzed using the thermal-optical transmission (TOT) technique with a Sunset carbon analyzer (Sunset Laboratory Inc., Portland, OR, USA) and the EUSAAR2 thermal protocol described in detail by Cavalli et al. [5].

The elemental composition of individual particles was assessed using single particle inductively coupled plasma time-of-flight mass spectrometry (SP-ICP-TOF-MS, TOFWERK, Thun, Switzerland), following protocols outlined in previous publications [6-8]. In brief, WFPM_0.1_ samples were introduced into the ICP with a 2DX autosampler (Element Scientific, Omaha, United States) and a MicroMist U-series Nebulizer (Thermo scientific, USA) connected via a Quartz Cyclonic Spray Chamber (Meinhard, USA) to the injector of the ICP torch. The instrument operating parameters and the monitored isotopes were published in [1].

PAHs samples from the peak combustion were quantified and analyzed using a modified protocol from Tsiodra et al., 2025 [9]. Briefly, WFPM_0.1_ suspensions were spiked with a known amount of deuterated PAHs (16 members) mixture (CPA Chem) before undergoing pressurized liquid extraction (PLE) using a 50:50 n-hexane -dichloromethane solvent mix. Extracts were purified via silica column chromatography, and the PAH-containing fraction was concentrated. The fraction containing PAHs was isolated by eluting with 11 mL of n-hexane and ethyl acetate mixture (8:2, v/v), then concentrated to a final volume of 0.1 mL. Before storage, a precise amount of [²H₁₂]perylene was added as an internal standard. Quantification was done using gas chromatography/mass spectrometry (GC/MS) (Agilent 7890/5975C) with calibration based on native and deuterated PAH mixtures to calculate the relative response factors (RRFs). Background contamination was assessed using blank quartz filters.

**Dispersion and colloidal characterization of** WFPM_0.1_ **suspensions**

Preparation and colloidal characterization of LS- and C-WFPM dispersions were carried out as we previously described [10-12]. Briefly, stock dispersions were prepared at 1 mg/mL in HyPure endotoxin-free water (HyClone, Cytiva, USA). Each suspension was subjected to one-minute rounds of cup-horn sonication (Branson Sonifier S–450D, 400 W, 3-inch horn; output 1.26 W), followed by 30 seconds of vertexing. After each sonication cycle, the hydrodynamic diameter (z-average, dH) was determined by dynamic light scattering (DLS; Zetasizer Nano ZS, Malvern Panalytical, Westborough, MA). Sonication rounds were continued until sequential measurements showed less than a 5% reduction in dH. The cumulative sonication duration and instrument power were then used to calculate the critical delivered sonication energy (DSEcr, J/mL) for each particle type. Final 1 mg/mL aqueous suspensions of C- WFPM_0.1_ and LS- WFPM_0.1_ were dispersed to their respective DSEcr values, followed by dilution in complete culture medium (used for all in vitro studies) to generate working treatment concentrations for the toxicological assays.

**Endotoxin and microbiological sterility testing of WFPM_0.1_ suspensions**

Endotoxin levels in C- WFPM_0.1_ and LS- WFPM_0.1_ suspensions were evaluated using the HEK-Blue™ LPS Detection Kit (Invivogen, San Diego, CA, USA) in accordance with the manufacturer’s protocol. Test wells included WFPM_0.1_ suspensions (100 µg/mL), extraction controls, and endotoxin-free water, alongside an *E. coli* O55 endotoxin standard curve (0.01–1 EU/mL). HEK-Blue™-4 cells were incubated with test samples for 20 h, followed by colorimetric detection with Quanti-Blue™ reagent at 620 nm. To confirm assay performance, selected C- WFPM_0.1_ and LS- WFPM_0.1_ suspensions were spiked with 0.1 EU/mL endotoxin.

Sterility of C- WFPM_0.1_ and LS- WFPM_0.1_ and vehicle control were verified following WHO pharmacopoeia guidelines [3, 4]. Suspensions (1 mg/mL) were cultured in thioglycolate medium for 14 days at 37 °C and inspected daily for contamination. Parallel plating on potato dextrose agar (PDA) and plate count agar (PCA) was performed to detect bacterial and fungal colonies throughout the incubation period.

**Biochemical analysis in BAL fluid**

**Evaluation of cell membrane integrity (LDH release)**

Collected BAL samples were centrifuged at 3,000 × g for 5 minutes to remove cells and debris. Fifty microliters of the clarified supernatant from each sample were transferred in triplicate to a 96-well plate and mixed with 50 µL of the LDH reaction mixture prepared according to the CyQUANT LDH Cytotoxicity Assay protocol (Thermo Fisher, Waltham, MA). Plates were incubated at room temperature for 30 minutes, and the reaction was stopped with 50 µL of stop solution. Absorbance was measured at 490 nm (A490) and 680 nm (A680), with background correction applied by subtracting A680 from A490. Percent cytotoxicity was calculated by subtracting background-corrected spontaneous LDH release values from background-corrected treatment values, dividing by total LDH activity (Maximum LDH release – Spontaneous LDH release), and multiplying by 100.

Cellular membrane integrity was assessed by measuring lactate dehydrogenase (LDH) levels released to the culture medium, using the CyQUANT LDH Cytotoxicity Assay (Thermo Fisher, Waltham, MA) according to the manufacturer’s instructions. Briefly, LDH substrate provide in the kit was dissolved in 11.4 mL of provided ultrapure water and added to 0.6 mL provided assay buffer to prepare the assay reaction mixture. Following 24-hour exposure of adherent PMA differentiated macrophages to HDPE-I, media alone (untreated – Spontaneous LDH release), or 45 min incubation with provided lysis buffer (positive control – Maximum LDH release), cell media from wells was collected in 1.5 mL tubes, and centrifuged at 3000 × g for 5 minutes to pellet cell debris. Fifty μL of the supernatant from each tube was dispensed in triplicate wells of a new 96-well plate along with 50 μL of the reaction mixture. Plates were then incubated at room temperature for 30 minutes before 50 mL of stop solution was added to each well to end the reaction. Absorbance was measured at 490 nm (A_490_) and 680 nm (A_680_) using a SpectraMax M-5 reader and SoftMax Pro software (Molecular Devices). A680 values were subtracted from A490 values to correct for instrument background. Percent cytotoxicity was calculated by subtracting background-corrected spontaneous LDH release values from background-corrected treatment values, dividing by total LDH activity (Maximum LDH release – Spontaneous LDH release), and multiplying by 100.

**Results**

**Endotoxin and sterility analysis**

Following a 14-day incubation, agar plates exposed to LS- and C- WFPM_0.1_ and control vehicle aqueous suspensions showed no microbial growth. Additionally, endotoxin assessment of all samples revealed levels below the assay’s detection threshold of 0.0079 EU/mL, indicating the absence of detectable endotoxin.

**Characterization of C-WFPM_0.1_ and LS-WFPM_0.1_ colloidal properties in water and culture media**

Results of colloidal properties of C- WFPM_0.1_ and LS- WFPM_0.1_ suspensions are presented in **Table S12**. In water, C- WFPM_0.1_ displayed a slightly larger hydrodynamic diameter (284.7 ± 71.4 nm) and higher polydispersity (PdI = 0.368 ± 0.007) compared to LS- WFPM_0.1_ (240.0 ± 4.6 nm, PdI = 0.103 ± 0.023). Both C-WFPM (−34.3 ± 0.3 mV) and LS- WFPM_0.1_ (−27.4 ± 11.8 mV) carried strongly negative surface charges, demonstrating an effective electrostatic stabilization of the particles and a favorable colloidal dispersion.

In complete medium, C- WFPM_0.1_ exhibited slightly larger particle size (314.5 ± 9.0 nm) relative to LS- WFPM_0.1_ (292.2 ± 1.3 nm), though both maintained similar polydispersity values (0.272 and 0.266, respectively). Additionally, both particle types showed a more positive zeta potential (C-WFPM: −8.14 ± 0.44 mV; LS- WFPM_0.1_: −9.86 ± 0.70 mV) compared to particles in aqueous suspensions. This is consistent with protein corona formation and charge neutralization.

Furthermore, conductivity increased substantially for both suspensions in media compared, with C- WFPM_0.1_ (14.8 ± 0.86 mS/cm) slightly higher than LS- WFPM_0.1_ (13.2 ± 1.01 mS/cm).
